# Inhibiting Clarinet/CLA-1 restores function to injured motor neurons

**DOI:** 10.1101/2025.07.05.663315

**Authors:** Wenjia Huang, Eli Min, Alexandra B. Byrne

**Affiliations:** Department of Neurobiology, University of Massachusetts Chan Medical School, Worcester, MA, 01605, USA; NeuroNexus Institute, University of Massachusetts Chan Medical School, Worcester, MA, 01605, USA; Interdisciplinary Graduate Program, University of Massachusetts Chan Medical School, Worcester, MA, 01605, USA; Medical Scientist Training Program, University of Massachusetts Chan Medical School, Worcester, MA, 01605, USA

## Abstract

To regain function, injured axons need to both regenerate and reform synapses with appropriate postsynaptic cells. We found that inhibiting the scaffolding protein Clarinet/CLA-1, a *C. elegans* ortholog of Piccolo and Fife, robustly improves axon regeneration. Despite the importance of CLA-1 during synapse development, disrupting the medium isoform of CLA-1 increases the number of axons that regenerate to the neuromuscular junction without significantly influencing synapse reformation. Consequently, the axons that do regenerate are capable of regaining function. Mechanistically, the enhanced axon regeneration observed in *cla-1(-)* mutants depends on the function of PTRN-1, a microtubule minus-end binding protein. Our data supports a model where loss of CLA-1 promotes PTRN-1 function, which speeds trafficking of injury-related cargo to and from the lesion, thus improving repair. Together, our results reveal a highly conserved synaptic active zone protein that can be manipulated to enhance axon regeneration without sacrificing the function of the repaired axons.

**Highlights:**

- Clarinet/CLA-1 is a robust inhibitor of *C. elegans* GABAergic axon regeneration
- Loss of CLA-1 function improves both axon regeneration and functional repair
- CLA-1 isoforms differentially regulate axon regeneration and synaptic transmission
- CLA-1 regulates axon regeneration via microtubule minus-end protein PTRN-1/Patronin
- CLA-1 knockdown improves cargo trafficking during the early injury response

## Introduction

Axonal injury severs neurons from their interacting cells and disrupts their function. Injured axons in the mammalian central nervous system (CNS) are unable to regenerate and others lose their ability to regenerate with age, resulting in permanent loss of sensory and motor function. In contrast, injured axons in many invertebrates and in the young mammalian peripheral nervous system are capable of regeneration, demonstrating this form of repair is feasible. Developing a comprehensive molecular and cellular understanding of how axon regeneration is positively and negatively regulated in axons that are capable of repair, as well as those that are not, is critical to developing approaches to manipulate the injury response towards repair.

*C. elegans* is a powerful model organism to screen for novel regulators of axon regeneration and determine their molecular and cellular mechanisms of action. Approximately 83% of *C. elegans* genes and 75% of human disease genes are conserved in these two species (Lai et al. 2000; Silverman et al. 2009; Apostolakou et al. 2021). This genetic conservation includes critical promoters and inhibitors of axon regeneration, such as the DLK-1/DLK/LZK mitogen-activated protein (MAP) kinase cascade and DAF-18/ phosphatase and tensin homologue (PTEN), which are conserved among *C. elegans*, *Drosophila* and mammals (Hammarlund et al. 2009; Yan et al. 2009; Xiong et al. 2010; Shin et al. 2012; Park et al. 2008; Liu et al. 2010; Song et al. 2012; Byrne et al. 2014). In addition to its highly conserved genome, each neuron in the *C. elegans* nervous system is well characterized and can be visualized *in vivo* through the animal’s transparent cuticle. These characteristics facilitate identifying and investigating conserved regulators of the injury response with single axon resolution (Yanik et al. 2004; Wu et al. 2007; Gabel et al. 2008; Ghosh-Roy et al. 2010; Chen et al. 2011; Nix et al. 2014; Kim et al. 2018; Czech et al. 2023).

To regain function, injured axons not only need to regenerate towards their postsynaptic targets, they must also reform synapses with the appropriate interacting cells, a process called functional axon regeneration. While regenerated axons are capable of reforming synapses and rewiring into proper circuits, degrees of restored neuronal function are limited and variable depending on the age and neuron-type, suggesting functional recovery is not a spontaneous consequence of axon regeneration (Yanik et al. 2004; El Bejjani and Hammarlund 2012; Byrne et al. 2016; Ding and Hammarlund 2018; Belew et al. 2023).

Here, we describe our findings that disrupting the function of active zone scaffolding protein CLA-1 robustly improves functional axon regeneration. CLA-1 and its orthologs Piccolo and Fife are scaffolding proteins in synaptic active zones that regulate synaptic vesicle clustering and release as well as autophagy (Xuan et al. 2017; Xuan et al. 2023; Krout et al. 2023). From a genetic screen, we found that injured motor axons lacking CLA-1 regenerate more often and further than wild-type motor axons. Despite the significant role of CLA-1 in active zone organization and function during development, loss of CLA-1 function in injured axons not only enhances regeneration, it also significantly improves synaptic vesicle trafficking and motor function relative to wild-type. Enhanced axon regeneration in *cla-1* mutant animals requires PTRN-1, a highly conserved microtubule minus-end binding protein homologous to the verterbrate calmodulin-regulated spectrin-associated protein family(CAMSAP) and Patronin in *D. melanogaster*, to regulate the speed and direction of cargo trafficking during the early injury response. Subsequent genetic epistasis analysis between CLA-1 and other active zone proteins reveals they regulate axon regeneration through multiple independent mechanisms. Together, our data reveal a previously uncharacterized inhibitor of axon regeneration, CLA-1/Piccolo/Fife, which functions cell-intrinsically with the microtubule minus-end binding protein PTRN-1 to regulate microtubule transport dynamics and the regenerative capacity of injured axons.

## Results

### Candidate screen reveals scaffolding protein Clarinet/CLA-1 is a robust inhibitor of axon regeneration

We screened candidate regulators of axon regeneration by severing individual fluorescently labeled GABAergic motor commissures at their midpoint with a pulsed nitrogen laser, and quantifying regeneration as the proportion of injured axons that form growth cones (Figure 1A). We found that Clarinet/CLA-1, a recently discovered active zone cytomatrix scaffolding protein that is highly homologous to mammalian Piccolo and Bassoon, inhibits repair. GABA motor axons lacking CLA-1 regenerated more frequently compared to wild-type GABA motor axons. 24 hours after injury, 76.4% of injured axons in *cla-1(ok2285)* null mutants formed regenerative growth cones compared to 61.4% in wild-type animals (p=0.02, Fisher’s exact test) (Figure 1D). In addition, injured axons lacking CLA-1 regenerated further than injured wild type axons. *cla-1(ok2285)* mutants extended 65.7% of their original length compared to 52.3% in wild-type animals (p=0.0017, Kruskal-Wallis test with Dunn’s multiple comparisons test) (Figure 1E). Of note, *cla-1(ok2285)* axons had a relatively wild-type morphology before injury, suggesting loss of CLA-1 function does not disrupt developmental axon outgrowth. Together, our results reveal CLA-1 as a robust inhibitor of axon regeneration.

**Figure 1.**
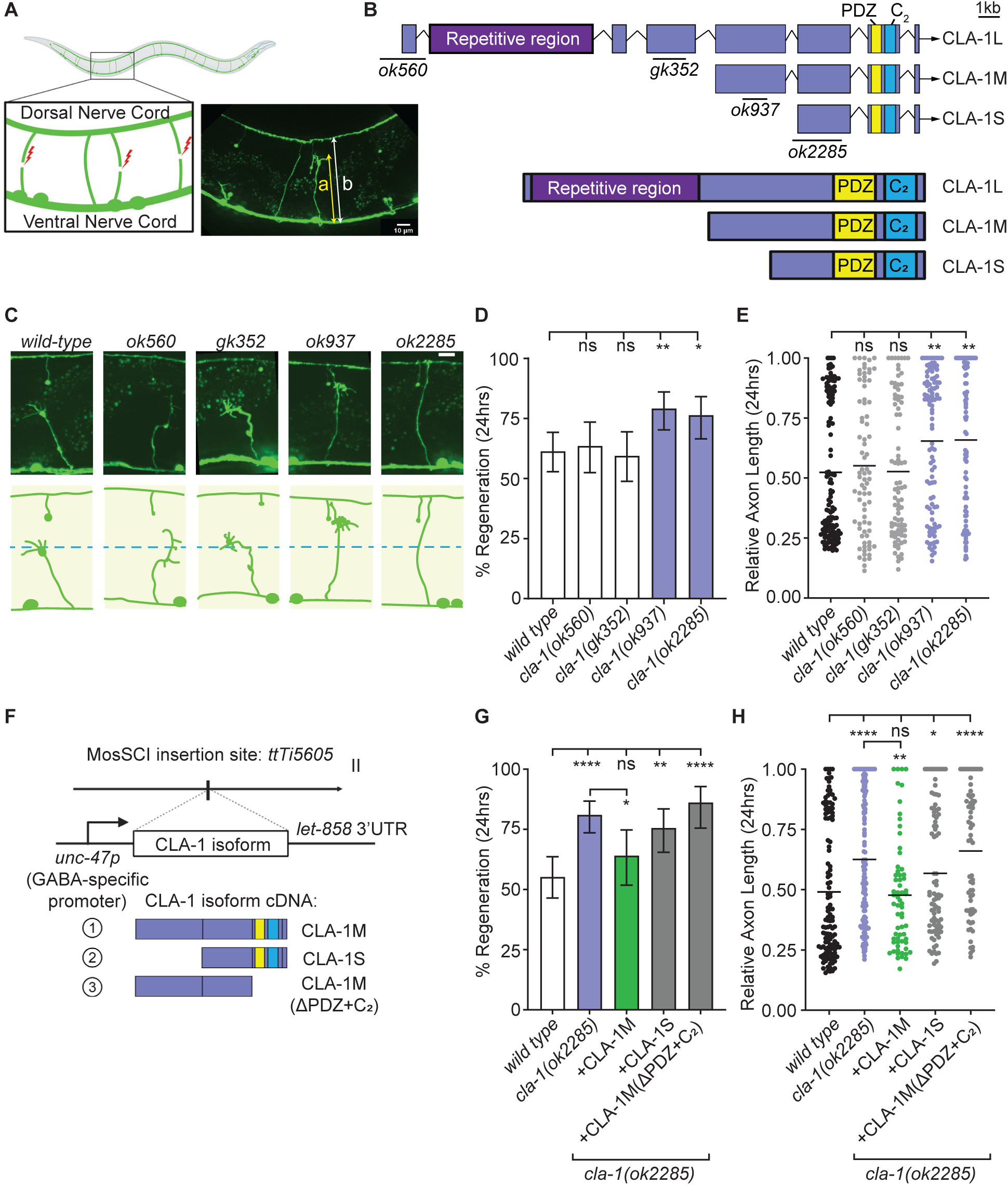
Clarinet/CLA-1 inhibits axon regeneration cell-autonomously through its medium isoform. (**A**) We axotomized GFP labeled GABA motor neurons in L4 animals and the dorsal-ventral midline and quantified regeneration 24 hours later by calculating the percentage of injured axons that initiated regeneration, as well as the distance axons regenerated, indicated by the yellow double-headed arrow, normalized to the width of animals, indicated by the white double-headed arrow. (**B**) Cartoon schematics of CLA-1 genetic locus which encodes three major isoforms: long (CLA-1L), medium (CLA-1M), and short (CLA-1S). Deletion alleles of *cla-1*, *ok560, gk352, ok937* and *ok2285*, affect different isoforms of CLA-1. (**C**) Representative regenerating axons in wild-type, *cla-1(ok560), cla-1(gk352), cla-1(ok937)*, and *cla-1(ok2285)* animals. Blue dashed line represents the dorsal-ventral midline. Scale bar: 10 µm. (**D and E**) Deletion alleles *ok560* and *gk352* that disrupt N-terminal of CLA-1L isoform do not affect axon regeneration. Injured axons in *cla-1(ok937)* animals that contain an in-frame deletion in CLA-1L and CLA-1M isoforms but not in the CLA-1S isoform, and in *cla-1(ok2285)* putative null animals regenerate more frequently and further 24 hours after injury in comparison to wild-type animals. (**D**) A higher percentage of injured axons in *cla-1(ok937)* and *cla-1(ok2285)* animals, but not in *cla-1(ok560)* and *cla-1(gk352)* animals, initiate regeneration compared to wild-type animals 24 hours after injury. N = 132, 77, 85, 101, 89. Significance relative to wild type is indicated by *P < 0.05, **P < 0.01, Fisher exact test. Error bars indicate 95% confidence intervals. (**E**) Injured axons in *cla-1(ok937)* and *cla-1(ok2285)* animals, but not in *cla-1(ok560)* and *cla-1(gk352)* animals, also regrow further along the dorsal-ventral axis 24 hours after injury compared to wild type. N = 132, 77, 85, 101, 89. Significance relative to wild type is indicated by **P < 0.01, Kruskal-Wallis test with Dunn’s multiple comparisons test. Median is indicated by the black line. (**F**) Mos1-mediated Single Copy Insertion (MosSCI) strategy for GABA motor neuron-specific expression of CLA-1 isoform cDNA for CLA-1M, CLA-1S and CLA-1MΔPDZ+C_2_. (**G and H**) GABA-specific expression of CLA-1M, but not CLA-1S or CLA-1MΔPDZ+C_2_, robustly rescues the enhanced axon regeneration in *cla-1(ok2285)* animals back to wild-type level. N = 125, 137, 64, 86, 65. (**G**) Significance of frequency of regeneration relative to wild-type and *cla-1(ok2285)* null animals is indicated by *P < 0.05, **P < 0.01, ****P<0.0001, Fisher exact test. Error bars indicate 95% confidence intervals. (**H**) Significance of regenerating axon length compared across wild-type and *cla-1(ok2285)* animals is indicated by *P < 0.05, **P < 0.01, ****P<0.0001, Kruskal-Wallis test with Dunn’s multiple comparisons test. Black bar represents median.

### CLA-1 inhibits axon regeneration cell-autonomously through its central uncharacterized region and C-terminal PDZ and C_2_ domains

CLA-1 and its orthologs Piccolo, Bassoon, and Fife are scaffolding proteins that are enriched at active zones (Xuan et al. 2017). *C. elegans* CLA-1 contains a unique N-terminal repetitive domain, a central uncharacterized region, and C-terminal PDZ and C_2_ domains, which are all highly conserved between CLA-1 and its orthologs (Figure S1). The *cla-1* gene encodes three major isoforms, categorized by their relative protein sizes as CLA-1L (long, ∼9000 amino acids), CLA-1M (medium, ∼2700amino acids) and CLA-1S (short, ∼1300 amino acids) (Figure 1B).

To understand which of the CLA-1 isoforms is responsible for inhibiting axon regeneration, we quantified regeneration in three additional deletion alleles that differentially disrupt the CLA-1 isoforms. Similar to *cla-1(ok2285)* null mutants, we observed a robust enhancement of axon regeneration in *cla-1(ok937)* animals harboring an in-frame deletion that disrupts the coding region of CLA-1L and CLA-1M, but not CLA-1S (Figure 1C-E). In addition, we tested two different alleles, *ok560* and *gk352*, that specifically affect CLA-1L or the N-terminus of CLA-1, but we did not observe a significant defect in axon regeneration (Figure 1C-E). Our finding that targeting CLA-1L and CLA-1M, but not CLA-1L alone, enhances axon regeneration to a similar degree as loss of all CLA-1 isoforms and raises a possibility that CLA-1M isoform plays a selective role in the injury context to inhibit axon regeneration.

To further test a distinct function of CLA-1M during axon regeneration, we took advantage of the MosSCI (Mos1-mediated Single Copy Insertion) genomic editing protocol to transgenically express an integrated single copy of CLA-1M cDNA driven by a GABAergic motor neuron-specific promoter *(unc-47p*) in *cla-1(ok2285)* null mutant animals (Figure 1F) (Frøkjaer-Jensen et al. 2008). We observed that expression of CLA-1M in GABAergic motor neurons was sufficient to rescue the frequency of axon regeneration and length of regenerating axons in injured *cla-1(ok2285)* mutants back to wild-type levels (Figure 1G,H). These data support the allelic-series results that CLA-1M has a unique inhibitory function relative to the other CLA-1 isoforms and that it functions cell-autonomously to inhibit axon regeneration. We did not observe rescue when we expressed CLA-1S or CLA-1M(ΔPDZ+C_2_), a C-terminal truncated CLA-1M that lacks highly conserved PDZ and C_2_ domains, in *cla-1(ok2285)* mutants (Figure 1G,H). We conclude that both the central uncharacterized region and highly conserved C-terminal PDZ and C_2_ domains of CLA-1M are required, while each component alone is not sufficient, for its ability to inhibit axon regeneration. Together, our findings indicate the medium isoform of CLA-1 functions cell-intrinsically to inhibit motor axon regeneration *in vivo*.

### CLA-1 functions in parallel to UNC-13, a core vesicle priming protein, to inhibit axon regeneration

Active zone cytomatrix proteins form intricate protein-protein interactions that support synapse organization and synaptic vesicle release (Figure 2A). To determine whether active zone function inhibits axon regeneration, we asked whether the amount of disruption a mutation confers on active zone organization and synaptic activity correlates with enhanced regeneration. We tested a panel of active zone mutations with increasing levels of locomotion defects. We first tested another active zone scaffolding protein ELKS-1, the sole *C. elegans* homolog of mammalian ELKS proteins. *elks-1 (ok2762)* null mutants move phenotypically wild-type, suggesting its minor role in synaptic transmission. In addition, active zones in *elks-1 (ok2762)* null mutants also display normal dense projection ultrastructure (Kittelmann et al. 2013) and wild-type expression of calcium channel UNC-2 (Oh et al. 2021). However, in contrast to the enhanced regeneration in *cla-1(ok2285)* mutants, we did not detect significant axon regeneration defects in *elks-1(ok2762)* mutants. These data indicate inhibition of axon regeneration is not a shared property of active zone scaffolding proteins (Figure 2B,C).

**Figure 2.**
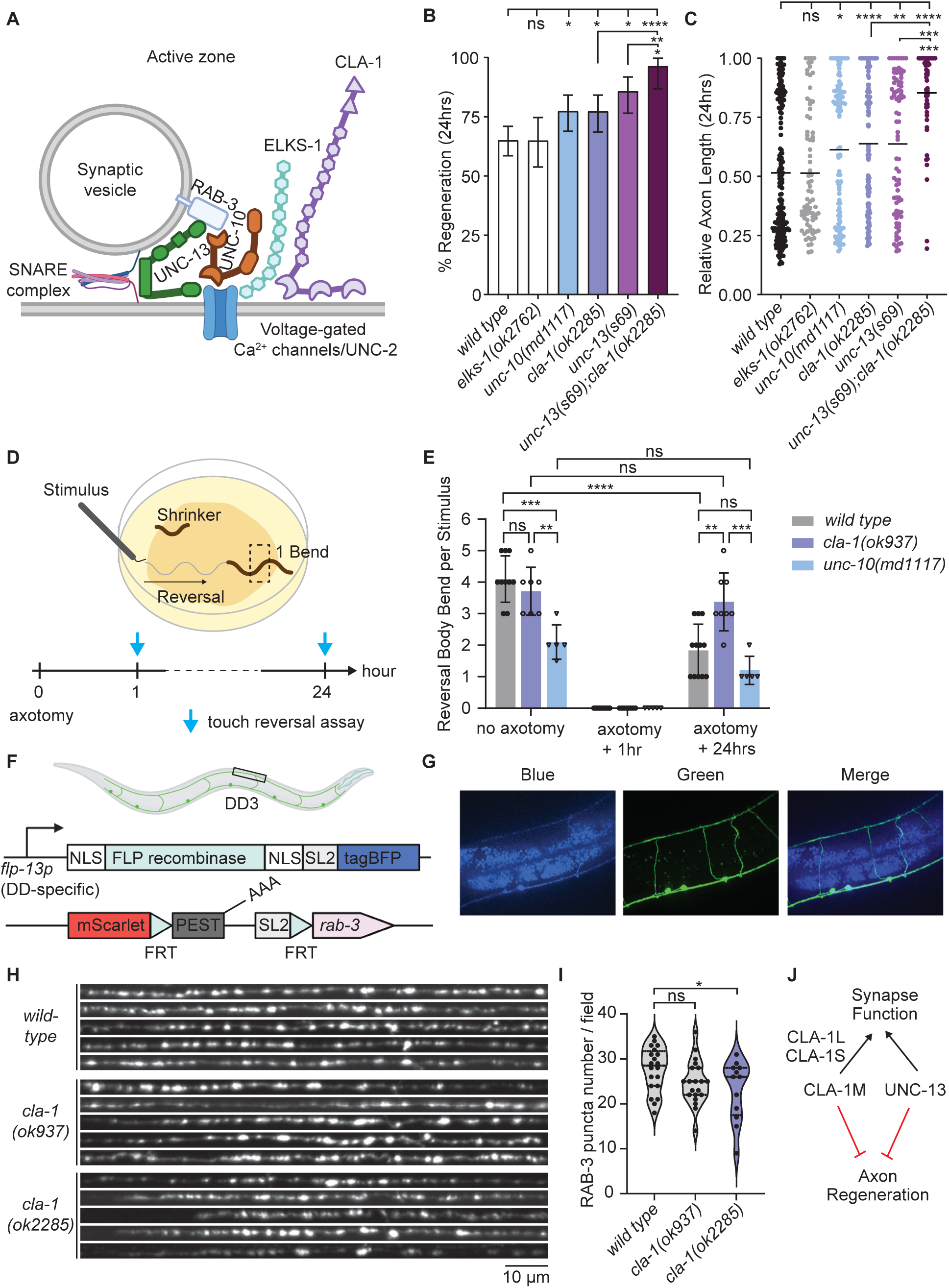
CLA-1 inhibits axon regeneration independently of its synaptic function at the active zone. (**A**) Simplified cartoon of *C. elegans* active zone (AZ). *C. elegans* AZs are specialized structures for neurotransmitter release upon Ca^2+^ influx through voltage-gated Ca^2+^ channels UNC-2/CaV2. Uncoordinated-10 (UNC-10)/Rab3-interacting molecule (RIM), a core AZ protein, binds to SV-associated RAB-3/Rab3 and prime SVs for release with other AZ proteins including UNC-13/Munc13, ELKS-1/protein rich in the amino acids E, L, K, and S (ELKS), CLA-1/Piccolo/Bassoon and SNARE complex. (**B and C**) Severing individual GABA neurons in *wild-type*, *elks-1(ok2762)*, *unc-10(md1117), cla-1(ok2285)*, *unc-13(s69)* and *cla-1(ok2285);unc-13(s69)* mutants indicates inhibiting axon regeneration is not a shared property of active zone proteins and is not correlated with the magnitude of their contributions to synaptic function. In addition, CLA-1 inhibits axon regeneration independently of UNC-13. Therefore, there are multiple mechanisms that active zone proteins utilize to inhibit axon regeneration. N = 223, 77, 115, 110, 84, 54. (**B**) Significance of frequency of regeneration relative to wild-type and *unc-13(s69);cla-1(ok2285)* animals is indicated by *P < 0.05, **P < 0.01, ****P<0.0001, Fisher exact test. Error bars indicate 95% confidence intervals. (**C**) Significance regenerating axon length relative to wild-type and *unc-13(s69);cla-1(ok2285)* animals is indicated by *P < 0.05, **P < 0.01, ***P<0.001, ****P<0.0001, Kruskal-Wallis test with Dunn’s multiple comparisons test. Black bar represents median. (**D**) Cartoon illustration of nose-touch reversal assay. To assess functional axon regeneration, all GABA motor commissures, excluding one proximal to the head, are injured in individual animals. The number of reverse body bends upon nose-touch stimulus are measured in axotomized animals and sham controls. (**E**) One-hour post-injury, axotomized animals are not capable of reversing upon nose-touch regardless of their genotype. 24 hours after injury when regenerated axons form functional connections, *cla-1(ok937)* animals recover significantly more motor function compared to wild-type and *unc-10(md1117) null* animals. Without injury, *cla-1(ok937)* mutants have a wild-type reversal response upon nose-touch stimulus in contrast to *unc-10(md1117)* mutants, suggesting CLA-1 plays a minor role compared to UNC-10 during synaptic transmission in *C. elegans* motor neurons. N= 10, 7, 5, 12, 8, 5, 12, 8, 5. Two-way ANOVA with Šídák’s multiple comparisons test is used to determine statistical significance. **P < 0.01, ***P<0.001, ****P<0.0001. (**F**) Cartoon illustration of tissue-specific recombination strategy to visualize endogenous SV-associated protein RAB-3. (**G**) FLP recombinase is highly expressed in DD neurons. However, weak leaky expression of the FLP recombinase is also observed in VD cell bodies, but not in other neuron types. Synapses (boxed region on the dorsal nerve cord) labeled by FLP-on mScarlet::RAB-3 were examined. (**H**) Straightened synaptic region of interest indicated in **F** showing the localization of FLP-on mScarlet::RAB-3 in wild-type, *cla-1(ok937)* and *cla-1(ok2285)* animals. (**I**) RAB-3 puncta number is reduced in *cla-1(ok2285)* mutants compared to wild-type controls, but not in *cla-1(937)* mutants. N=20, 20, 12. One-way ANOVA with Dunnett’s multiple comparisons test. *P < 0.05. (**J**) CLA-1 isoforms have differential roles in regulating synaptic function and axon regeneration.

We next investigated axon regeneration in mutants whose active zones are significantly disrupted. Null mutations in *unc-10/RIM* and *unc-13/MUNC13*, which are critical for priming vesicle release, cause significantly more severe locomotion defects compared to *cla-1(ok2285)* animals: loss of *unc-10(md1117)* function results in increased body bends (Koushika et al. 2001) and loss of *unc-13(s69)* function results in paralysis (Richmond, Davis, and Jorgensen 1999). In mammals, conditional knockdown of RIM and MUNC13 enhances axon outgrowth of injured murine dorsal root ganglion (DRG) neurons *in vitro*, and loss of MUNC13 enhances axon regeneration *in vivo* in a spinal cord injury model (Hilton et al. 2022). Consistent with these mammalian findings, we observed a robust increase in GABAergic motor axon regeneration in *unc-10(md1117)* and *unc-13(s69)* mutants (Figure 2B,C). In conclusion, the amount of synaptic function that is disrupted by *cla-1(ok2285), elks-1(ok2762), unc-10(md1117)* or *unc-13(s69)* single mutation differs significantly; however, we observed a similar increase in axon regeneration in *cla-1(ok2285), unc-10(md1117)* and *unc-13(s69)* single mutants. These data indicate that the ability of CLA-1 to inhibit axon regeneration is not correlated with the magnitude of its contribution to synaptic function (Figure 2B,C).

Because loss of CLA-1 and UNC-10 leads to a synergistic reduction in UNC-13 expression and synaptic transmission at the neuromuscular junction (NMJ) (Xuan et al. 2017; Krout et al. 2023), we performed a genetic epistasis analysis to determine whether CLA-1 and UNC-13 function together to regulate axon regeneration. In *cla-1(ok2285);unc-13(s69)* double mutants, we observed further enhancement in axon regeneration compared to *cla-1(ok2285)* or *unc-13(s69)* single mutants (Figure 2B,C), indicating CLA-1 inhibits axon regeneration through a separated mechanism from its canonical active zone partner UNC-13, and that active zone proteins inhibit axon regeneration through multiple independent mechanisms.

### Injured *cla-1 loss-of-function* animals recover pre-injury level of motor function

To regain function, an injured axon must both regenerate and form a functional synapse, a process called functional axon regeneration. Our finding that CLA-1, a synapse protein, inhibits axon regeneration raises a theoretical challenge to promote axon regeneration and functional recovery at the same time. Although regeneration is improved upon loss of CLA-1, are those axons capable of forming functional synapses? To determine whether *cla-1(ok937)* animals are capable of functional axon regeneration, we first severed 15 of the 16 D-type GABA motor neuron axon commissures on the right side of the animal and measured their ability to regain motor function. The D-type GABA motor neurons are important regulators of sinusoidal locomotion. They form inhibitory synapses on body wall muscles along the length of the animal, while also sending rhythmic and coordinated signals to cholinergic axons that activate body wall muscles on the opposing side of the animal (White et al. 1986). Animals that lose D-neuron function simultaneously contract muscles on both sides and fail to elicit reversal response when touched on the head (McIntire et al. 1993). We quantified motor functions using the number of body bends animals made in response to a touch stimulus on their nose in wild-type, comparing *cla-1(ok937)* and *unc-10(md1117)* mutant animals before and after injury (Yanik et al. 2004; Byrne et al. 2016) (Figure 2D). We did not include *unc-13(s69)* animals because they are already paralyzed without injury. Consistent with the previously reported minor behavior defects of *cla-1* loss-of-function mutants (Xuan et al. 2017; Krout et al. 2023), we did not detect any baseline difference in the number of body bends per touch stimulus between *wild-type* and *cla-1(ok937)* animals before injury (Figure 2E). In contrast, we observed significantly reduced body bends in *unc-10(md1117)* mutants (Figure 2E). One hour after injury, before axons have initiated regeneration, axotomized *wild-type*, *cla-1(ok937)* and *unc-10(md1117)* animals exhibit “shrinker” phenotypes where they are unable to reverse in response to nose touch (Figure 2E). Twenty-four hours after injury, once axons have regenerated, axotomized *cla-1(ok937)* animals are capable of significantly more body bends compared to axotomized *wild-type* and *unc-10(md1117)* animals (Figure 2E). These results suggest that *cla-1(ok937)* mutants recover more motor function post-regeneration than wild type animals due to the increased number of axons that regenerate relative to wild type animals. In addition, *cla-1(ok937)* mutants recover more function than *unc-10(md1117)* animals because of the relatively minor disruption to synaptic transmission upon loss-of-function of *cla-1*.

We next examined the effect of *cla-1(ok937)* on synaptic vesicle distribution using a well-established synaptic vesicle marker RAB-3 GTPase (Figure 2F). To monitor endogenous RAB-3 expression in a tissue-specific manner, we took advantage of an established FLP/FRT-based recombination strategy to express flippase (FLP) recombinase under the *flp-13* promoter, which drives expression specifically in D-type GABA motor neurons (DD) (Kim and Li 2004; Davis et al. 2008). The DD-specific FLP recombinase was expressed in animals whose endogenous *rab-3* locus was previously modified with CRISPR-Cas9 to include an N-terminal FLP-ON::mScarlet cassette (Schwartz and Jorgensen 2016; McDonald, Fetter, and Shen 2020) (Figure 2F). To verify FLP recombinase expression, we added a trans-spliced blue fluorescent protein (SL2::BFP) reporter to the C-terminus of FLP recombinase. As expected, we observed high levels of BFP expression in DD motor neurons; however, we also noted very faint and leaky expression in the VD motor neurons that innervate the ventral body wall muscles, but we did not observe BFP expression in cholinergic motor neurons (Figure 2G). Using this FLP-ON strategy, we quantified the number of discrete RAB-3 labeled presynaptic terminal puncta along the dorsal nerve cord of DD neurons (Figure 2F). We observed a significant reduction of RAB-3 puncta number in *cla-1(ok2285)* null mutants compared to wild-type animals, consistent with the previous literature that loss of all CLA-1 isoforms causes a reduction in number of synapses (Figure 2H,I) (Xuan et al. 2017). We did not observe a significant decrease in the number of RAB-3 puncta number in *cla-1(ok937)* mutants (Figure 2H,I). Our intriguing finding that *ok937*, an in-frame deletion allele of CLA-1, does not affect synaptic vesicle development but robustly enhances axon regeneration suggests these two functions of CLA-1 may be decoupled at the isoform level, where CLA-1M plays a selective role to inhibit axon regeneration, and partial loss of CLA-1L and CLA-1M does not have a significant disruption on synapses (Figure 2J). Together, our results suggest the possibility to promote functional axon regeneration by targeting active zone scaffolding protein CLA-1. Specifically, targeting CLA-1 medium isoforms is a promising strategy to enhance axon regeneration without disrupting the developmental synaptic vesicles as well as post-axotomy functional recovery.

### PTRN-1 genetically suppresses the ability of CLA-1 to inhibit axon regeneration

Next, we sought to determine the molecular mechanism by which CLA-1 regulates axon regeneration. We performed a candidate screen of reported physical interactors of CLA-1 (candidates are listed in Table S1) (Lenfant et al. 2010; Sanchez et al. 2021; Artan et al. 2022), and found that loss of PTRN-1, a highly conserved minus-end microtubule binding protein homologous to mammalian CAMSAP, is required to mediate the inhibitory function of CLA-1 during axon regeneration (Figure 3A). In GABAergic motor neurons, loss of PTRN-1 by genetic deletion allele *tm5597* does not lead to any significant effect on axon regeneration, in contrast to the enhanced axon regeneration observed in *cla-1(ok937)* animals (Figure 3B-D). However, in *cla-1(ok937); ptrn-1(tm5597)* double mutants, we observed a reversal of *cla-1(ok937)* phenotype in axon regeneration (Figure 3B-D).This genetic interaction suggests that PTRN-1 function is required for the enhanced regeneration observed in *cla-1(ok937)* mutants.

**Figure 3.**
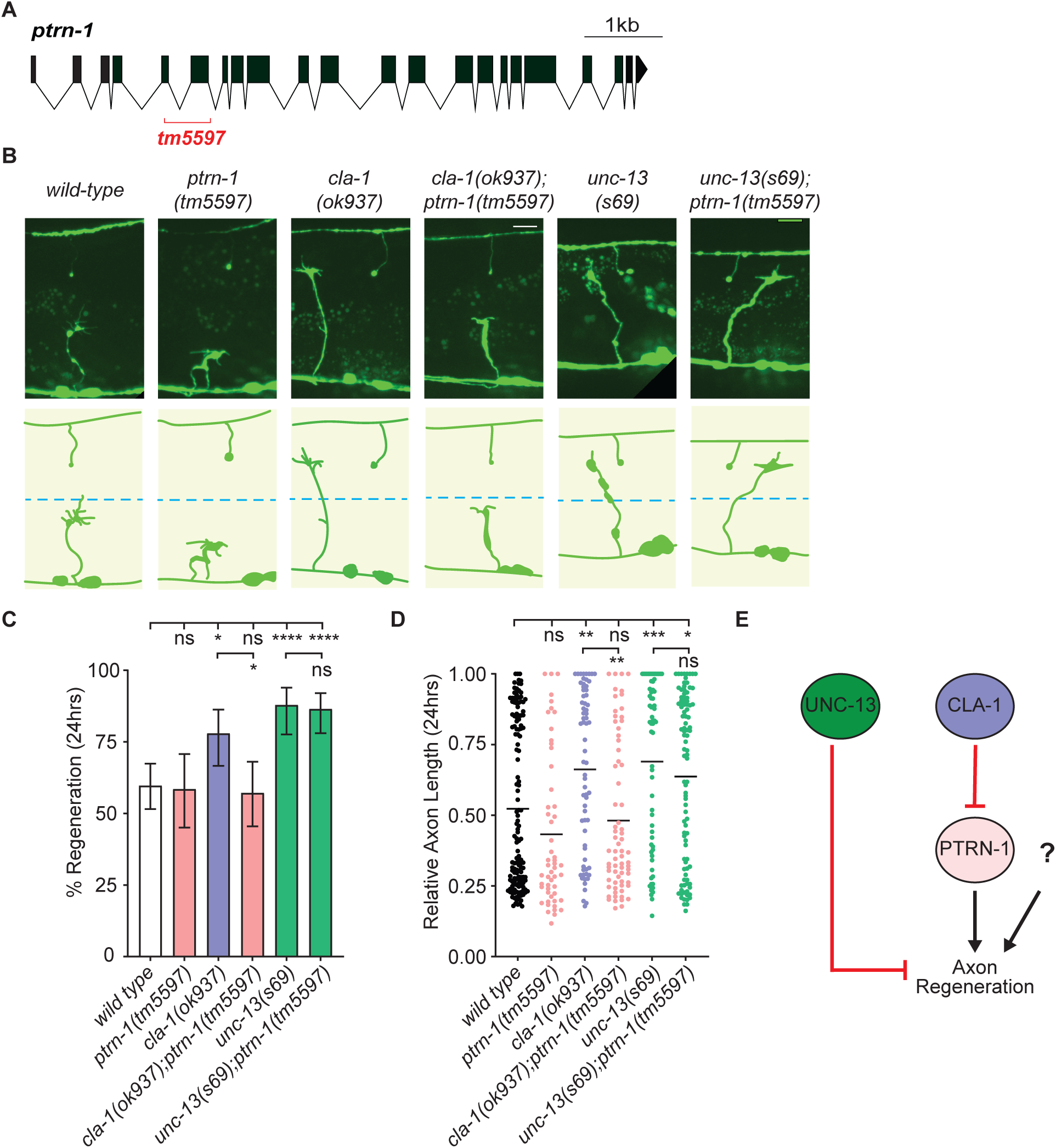
Microtubule minus-end binding protein PTRN-1/CAMSAP is required for CLA-1 to inhibit axon regeneration. (**A**) Schematic of *ptrn-1* genomic locus. *tm5597* is a putative null allele of *ptrn-1*. (**B**) Representative images of wild-type, *ptrn-1(tm5597)*, *cla-1(ok937), cla-1(ok937);ptrn-1(tm5597)*, *unc-13(s69), unc-13(s69);ptrn-1(tm5597)* injured axons. Scale bar: 10 µm. (**C and D**) Animals lacking PTRN-1 alone display wild-type axon regeneration. However, loss of PTRN-1 genetically suppresses the enhanced regeneration in *cla-1(ok937)* animals, but not the enhanced regeneration in *unc-13(s69)* animals. N = 144, 53, 68, 70, 66, 96. (**C**) Significance of frequency of regeneration relative to wild-type, *cla-1(ok937)* and *unc-13(s69)* animals is indicated by *P < 0.05, ****P<0.0001, Fisher exact test. Error bars indicate 95% confidence intervals. (**D**) Significance of regenerated axon length relative to wild-type *cla-1(ok937)* and *unc-13(s69)* animals is indicated by *P < 0.05, **P < 0.01, ***P < 0.001, Kruskal-Wallis test with Dunn’s multiple comparisons test. Black bar represents median. (**E**) CLA-1 and UNC-13 act in parallel to inhibit axon regeneration through different downstream mechanisms. PTRN-1 likely functions redundantly with other microtubule proteins to regulate axon regeneration, and PTRN-1 is required for CLA-1, but not UNC-13, to inhibit axon regeneration.

We next asked whether PTRN-1 is a universal suppressor of axon regeneration due to its important role in regulating microtubule structure and function in *C. elegans* neurons (Marcette, Chen, and Nonet 2014; Richardson et al. 2014; Chuang et al. 2014; He et al. 2022). We found that *unc-13(s69); ptrn-1(tm5597)* double mutants regenerated to the same extent as *unc-13(s69)* mutants (Figure 3B-D). The absence of suppression by the *ptrn-1* mutation on the enhanced axon regeneration in *unc-13(s69)* mutants (Figure 3B-D) indicates there is a specific genetic interaction between PTRN-1 and CLA-1 during axon regeneration, which further supports our finding that CLA-1 and UNC-13 function in parallel to inhibit axon regeneration using different downstream molecular mechanisms (Figure 3E).

Previously, PTRN-1 overexpression was reported to bypass the requirement for the mitogen-activated protein kinase signaling pathway that includes the MAPKKK DLK-1 and the transcription factor CEBP-1 (Chuang et al. 2014). Supporting our model that CLA-1 negatively regulates PTRN-1 during axon regeneration, we also observed an increase in regeneration in *cla-1(ok937);cebp-1(tm2807)* double mutants (Figure S2A-C). However, we did not observe a significant increase in growth cone formation in *dlk-1(ju476);cla-1(ok937)* double mutants. The requirement for DLK-1 may indicate that DLK-1 itself functions in parallel or downstream of CLA-1 (Figure S2A-C). Lastly, enhanced axon regeneration in *cla-1(ok937);cebp-1(tm2807)* double mutants is suppressed by loss of PTRN-1, furthering supporting our model that CLA-1 and PTRN-1 function in the same genetic pathway to regulate axon regeneration, at least partially independently, of CEBP-1 (Figure S2A-C).

### CLA-1 is not observed in the axon fragment proximal to the injury and is rapidly cleared from the distal fragment

An intriguing question that arises from our findings is how does CLA-1, a distal synaptic protein, regulate axon regeneration at the proximal side of the injury? One possibility is that CLA-1 is actively transported to the injury site following axotomy, and CLA-1 accumulation at the injury site may be deleterious for initiation of axon regeneration. To explore this possibility, we first monitored the endogenous CLA-1 expression with a CRISPR-tagged C-terminal GFP allele of CLA-1 (McDonald, Fetter, and Shen 2020). Consistent with previous reports (Xuan et al. 2017; McDonald, Fetter, and Shen 2020; Krout et al. 2023; Xuan et al. 2023), CLA-1::GFP is highly expressed in the nervous system with discrete puncta patterns that are likely to reflect the location of active zones (Figure 4A). To further monitor subcellular dynamics of CLA-1 expression in a tissue-specific manner, we used the FLP-FRT recombination system to activate CLA-1 expression in DD neurons (CLA-1::FLP-ON::mScarlet; Pflp-13::FLP::SL2::BFP) and characterized endogenous CLA-1 expression pattern prior to and 24 hours post injury (Figure 4B). We quantified CLA-1 puncta in DD commissures and dorsal neurites. We did not analyze CLA-1 signals on the ventral side of the animals as we cannot distinguish whether these CLA-1 signals are from DD neurons or neighboring VD neurons with some leaky FLP recombinase expression. Before injury, we observed CLA-1 signals along the dorsal nerve cord but not in the commissures of wild-type DD neurons (Figure 4C). After injury, we observe the loss of CLA-1 puncta along the dorsal nerve cord (Figure 4C,D). Loss of CLA-1 puncta in the injured distal stumps suggests possible synapse loss upon injury possibly due to activation of a degradation program upon axonal injury or normally present at the axonal neurites to maintain active zone protein dynamics. In contrast, at the proximal end of the injury, we did not detect any CLA-1 signals in regenerative growth cones or non-regenerative retraction bulbs, suggesting CLA-1 is not transported to the injured sites to block axon regeneration (Figure 4C). However, because we tagged the endogenous *cla-1* locus with FLP-ON::mScarlet cassette, it remains possible that CLA-1 may be present at the injured sites at an undetectable level.

**Figure 4.**
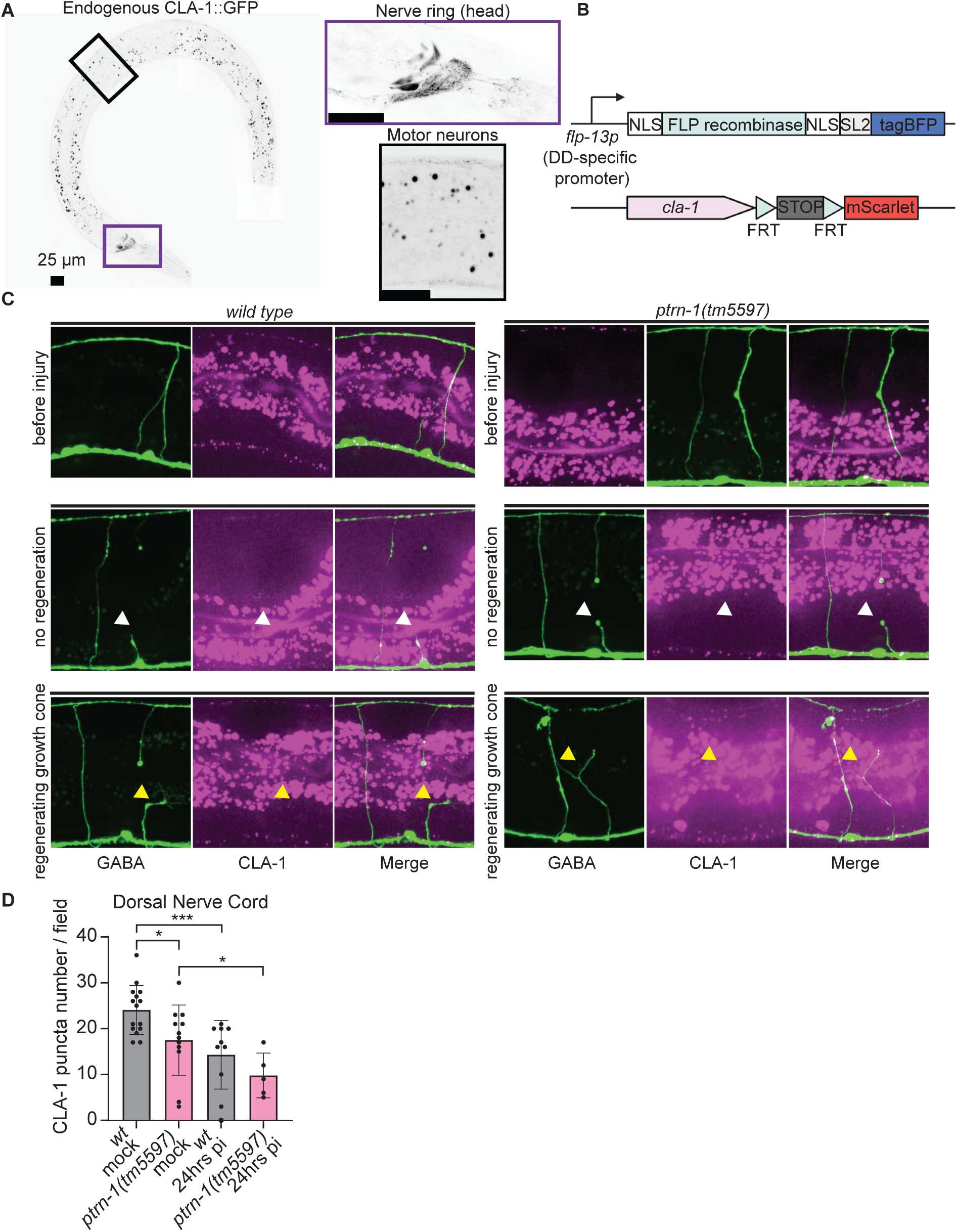
Axonal injury does not induce CLA-1 accumulation at the injury site. (**A**) Endogenous CLA-1::GFP expression is enriched in the *C. elegans* nervous system at the larval stage 4. (**B**) Tissue-specific recombination strategy to visualize endogenous CLA-1. (**C**) Green (GABA motor neuron), magenta (CLA-1::FLPon-mScarlet), and merged channels of uninjured, non-regenerative, and regenerating axons in wild-type and *ptrn-1(tm5597)* animals. (**D**) CLA-1 puncta number at the dorsal nerve cord is reduced in *ptrn-1(tm5597)* mutants before injury compared to wild-type controls. Axonal injury also leads to CLA-1 puncta reduction in both wild-type and *ptrn-1(tm5597)* animals as the distal stump is disconnected from the rest of the neuron. However, CLA-1 puncta signal was not observed in the cell bodies or in the proximal end of the injured axons in injured wild-type and *ptrn-1(tm5597)* animals. N=15, 12, 10, 5. Two-way ANOVA. *P < 0.05, ***P<0.001.

Given the robust genetic interaction between CLA-1 and PTRN-1 during axon regeneration, we next examined CLA-1 expression pattern in *ptrn-1(tm5597)* animals before and after axonal injury. Before injury, we observed a reduction of CLA-1 puncta number on the dorsal side, consistent with previous report that loss of PTRN-1 leads to aberrant expression of synaptic vesicle marker RAB-3 during development (Marcette, Chen, and Nonet 2014) (Figure 4C,D). We observed similar injury-induced changes in the CLA-1 expression pattern in the absence of PTRN-1: CLA-1 signals in the distal stump are further reduced in injured *ptrn-1(-)* animals, and there is also not a CLA-1 signal detectable at the regenerating growth cones or non-regenerating retraction bulbs (Figure 4C,D). Together, our results suggest CLA-1 is not recruited to the site of injury to physically block axon regeneration, but likely functions via an indirect mechanism through its interaction with PTRN*-*1. The requirement for PTRN-1 for wild type CLA-1 localization is only observed during development, but not during axon regeneration at the proximal side to mediate CLA-1 localization to the injury site, or at the distal side to modulate CLA-1 degradation in the stump.

### Cargo trafficking in *cla-1* mutants is faster during the early injury response

Animals lacking PTRN-1 display aberrant microtubule (MT) dynamics that are further exacerbated by axonal injury (Chuang et al. 2014). Destabilized MTs lead to functional defects in axonal cargo transport as neuronal microtubules are non-centrosomal arrays with partial overlaps, and this tiled organization of neuronal MTs requires cargo to constantly switch tracks at MT ends to traffic along the axon (Yogev et al. 2016). We therefore investigated whether the genetic interaction between PTRN-1 and CLA-1 reflected changes in microtubule functions by monitoring trafficking of endogenous RAB-3::FLP-ON mScarlet in DD commissures (Figure 5A). We observed that prior to injury RAB-3 puncta moves at a similar velocity in *wild-type* and *cla-1(ok937)* mutant animals, suggesting that loss of CLA-1 does not affect baseline RAB-3 trafficking speed (Figure 5B,C). Eight hours after injury, RAB-3 puncta just below the injury site moved at a slower velocity compared to mock controls in the *wild-type* group, likely to reflect the dynamic nature of damaged microtubules during the initial stage of axon regeneration (Figure 5B,C). Interestingly, commissural RAB-3 puncta in *cla-1(ok937)* animals maintain its pre-injury speed, significantly faster than injured *wild-type* animals (Figure 5B,C). We also examined the directionality of commissural RAB-3 trafficking. Before injury, RAB-3 puncta move both anterogradely and retrogradely at a similar frequency in *wild-type* and *cla-1(ok937)* animals. Eight hours after injury, we observed a significant bias toward retrograde movement in injured *wild-type* axons (Figure 5B,D). Significantly, such retrograde bias of RAB-3 trafficking was not detected in injured axons in *cla-1(ok937)* (Figure 5B,D). These data suggest that loss of CLA-1 exerts protection against injury-induced disruptions on microtubule function by preserving both the velocity and directionality of axonal cargo transport.

**Figure 5.**
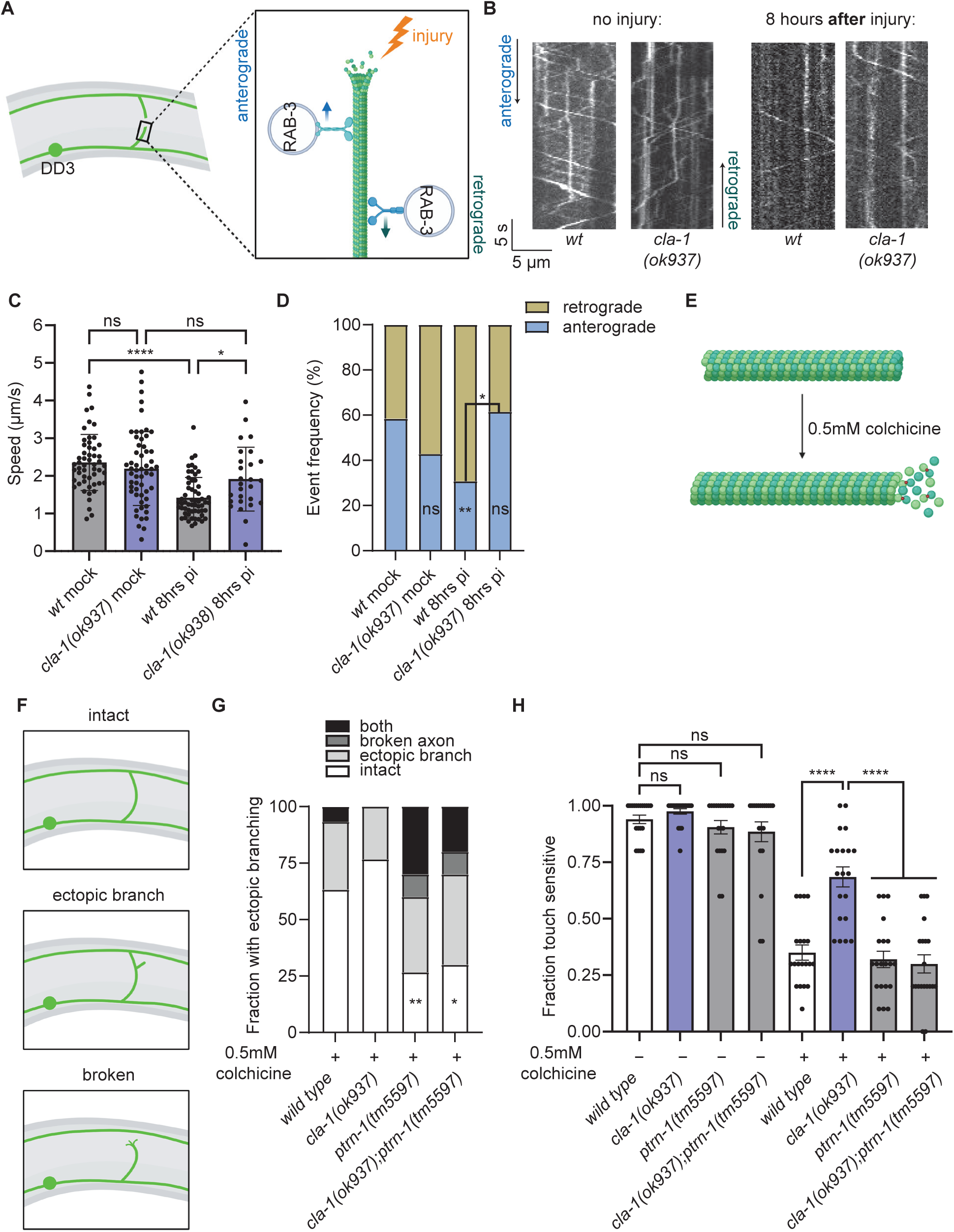
Loss of CLA-1 enhances microtubule stability and cargo trafficking in a PTRN-1-dependent manner. (**A**) Illustration of commissural FLP-on mScarlet::RAB-3 trafficking in injured DD axons. Commissural RAB-3 trafficking was recorded in the indicated region surrounding the site of injury for 30 seconds. (**B**) Representative kymographs of RAB-3 trafficking in wild-type and *cla-1(ok937)* animals before injury and 8 hours after injury. (**C**) RAB-3 puncta travel at reduced speeds in wild-type animals during the early injury response while they maintain their pre-injury speed in *cla-1(ok937)* animals. There is no difference in RAB-3 trafficking speed between wild-type and *cla-1(ok937)* animals before injury. N=53, 56, 52, 27. Two-way ANOVA with Šídák’s multiple comparisons test. *P < 0.05, ****P<0.0001. (**D**) RAB-3 puncta travel retrogradely at a higher frequency in wild-type animals during the early injury response while there is no directional bias in injured *cla-1(ok937)* animals. In addition, there is no difference in RAB-3 trafficking directionality between wild-type and *cla-1(ok937)* animals before injury. N=53, 56, 52, 27. Fisher exact test. **P < 0.01. (**E**) Colchicine (depicted as red dot) binds to alpha-and beta-tubulin monomers and chronic colchicine treatment induces microtubule depolymerization. (**F**) Illustration of aberrant morphologies, including ectopic branching and broken commissures, in wild-type, *ptrn-1(tm5597)*, *cla-1(ok937),* and *cla-1(ok937);ptrn-1(tm5597)* GABAergic motor neurons with 0.5mM colchicine treatment. (**G**) *ptrn-1(tm5597)* and *cla-1(ok937);ptrn-1(tm5597)* animals grown in 0.5mM colchicine display more severe morphological defects compared to wild-type controls, whereas *cla-1(ok937)* animals grown in 0.5mM colchicine display mostly intact GABAergic motor commissures. N=30, 30, 30, 30. Fisher exact test. *P < 0.05. (**H**) Under prolonged treatment of 0.5mM colchicine, *cla-1(ok937)* animals display higher sensitivity toward touch stimulus compared to wild-type animals. Touch sensitivity of *cla-1(ok937)* animals is abolished in *cla-1(ok937);ptrn-1(tm5597)* animals under 0.5mM colchicine treatment. There is no difference in touch sensitivity among wild-type, *ptrn-1(tm5597)*, *cla-1(ok937),* and *cla-1(ok937);ptrn-1(tm5597)* animals without colchicine treatment. N=20, 20, 20, 20, 20, 20, 20, 20. One-way ANOVA with Šídák’s multiple comparisons test. ****P<0.0001.

### Animals lacking CLA-1 are resistant to colchicine-induced microtubule disruption in a PTRN-1-dependent manner

To assess the role of CLA-1 in microtubule structure, we used colchicine to pharmacologically destabilize microtubules in *wild-type* and *cla-1(ok937)* animals and asked whether loss of CLA-1 also protects against colchicine-induced defects in microtubule structure. Colchicine binds both alpha-and beta-tubulin monomers to prevent microtubule polymerization (Steward, Goldschmidt, and Sutula 1984) (Figure 5E). In the presence of colchicine, *wild-type* neurons exhibit morphological defects, including ectopic branch and axon breakage, and functional defects in response to touch stimulus (Richardson et al. 2014) (Figure 5F). Consistent with the literature, we also observed ectopic branches and broken commissures in *wild-type* GABAergic neurons chronically treated with 0.5 mM colchicine (Figure 5G). The colchicine-induced defect in commissure morphology is further exacerbated upon loss of PTRN-1, supporting its reported role in stabilizing microtubules (Marcette, Chen, and Nonet 2014; Richardson et al. 2014). Significantly, *cla-1(ok937)* animals with chronic colchicine treatment display less ectopic branches and have no broken commissures (Figure 5G). However, improved structural integrity in *cla-1(ok937)* GABAergic commissures is abolished in *cla-1(ok937);ptrn-1(tm5597)* double mutants, suggesting PTRN-1 functions downstream of CLA-1 to maintain microtubule structure upon pharmacological insult (Figure 5G). At the behavioral level, we also found that animals lacking CLA-1 are more responsive to touch stimulus after chronic exposure to colchicine compared to *wild-type, ptrn-1(tm5597)* or *cla-1(ok937);ptrn-1(tm5597)* animals, while there is no significant difference among genotypes without colchicine treatment (Figure 5H). Together, these findings support a model where PTRN-1 functions antagonistically and downstream CLA-1 to modulate microtubule structure and function in response to pharmacological or physical insults.

## Discussion

The mechanisms that regulate functional axon regeneration, where injured axons must both regenerate and reform functional synapses, are poorly understood. Here, we find that knocking down the function of CLA-1/Piccolo improves both axon regeneration and functional recovery in *C. elegans* GABAergic motor neurons. CLA-1 and its highly conserved homologs, Piccolo and Bassoon, are active zone scaffolding proteins that promote synapse development. Therefore, our finding raises the perplexing question: how could eliminating the function of a synaptic protein improve functional repair? Our data indicate that disrupting the medium isoform of CLA-1 increases the number of axons that regenerate to the neuromuscular junction, without significantly influencing synapse development. Consequently, the axons that do regenerate are capable of regaining function. Mechanistically, the enhanced axon regeneration observed in *cla-1(-)* mutants depends on the function of PTRN-1, a microtubule minus-end binding protein. This molecular interaction led us to find that loss of CLA-1 function in injured axons upregulates the speed of cargo transport and reduces retrograde-bias during the early injury response. Our data support a model where CLA-1 inhibits PTRN-1 function, which impairs trafficking of injury-related cargo to and from the lesion, thus limiting repair. Together, our findings expand our understanding of how synaptic proteins inhibit axon regeneration and reveal a novel approach to improve functional repair.

### Disrupting CLA-1 improves axon regeneration without eliminating the synaptic function of regenerated axons

CLA-1 is a critical regulator of active zone structure and function during synapse development (Xuan et al. 2017; Krout et al. 2023). Despite its prominent role in synaptic active zone structure, animals lacking CLA-1 have superficially wild-type motor function, and electrophysiological recordings reveal synaptic transmission defects in *cla-1(-)* or *Piccolo^-/-^*animals are mild or not detectable compared to other active zone mutants, such as *unc-10(-)* animals(Mukherjee et al. 2010; Xuan et al. 2017; Krout et al. 2023).

Our findings genetically decouple the role of CLA-1 in axon regeneration from synaptic function with multiple lines of evidence. First, by quantifying axon regeneration in a panel of active zone mutants that are differentially required for synaptic function, including ELKS-1/ELKS, UNC-10/RIM and UNC-13/MUNC13, we found that regeneration frequency and distance does not correlate with the amount of synaptic function that is disrupted by each mutation (Figure 2B,C). For example, while null mutations in *cla-1* and *elks-1* have minor synaptic function defects compared to loss of function mutations in the other core active zone proteins, only *cla-1(-)* mutants display robust axon regeneration. On the other hand, while null mutations in *cla-1* and *unc-13* disrupt synaptic transmission to different degrees, *cla-1(-)* and *unc-13(-)* mutants have similar axon regeneration phenotypes. These data demonstrate the magnitude of synaptic function does not determine the propensity of an axon to regenerate. Second, we asked if enhanced axon regeneration correlates with defective synaptic vesicle expression and localization (RAB-3::FLP-ON mScarlet), which is a prominently used readout of synapse integrity (Schwartz and Jorgensen 2016; McDonald, Fetter, and Shen 2020). Our data indicate both a null mutation (*ok2285*) and a hypomorphic allele (*ok937*) of *cla-1* that introduces an in-frame deletion that only disrupts the coding sequence of the long and medium isoforms CLA-1L and CLA-1M, promote axon regeneration to a similar extent (Figure 1B,D,E) However, only the null allele significantly reduces the number of RAB-3 puncta in uninjured axons (Figure 2H,I). These data separate synaptic vesicle distribution from axon regeneration. Third, *cla-1(ok937)* animals regain more motor function compared to wild-type controls, suggesting regenerated axons in *cla-1(ok937)* animals are functional (Figure 2D). Together, our results indicate the inhibitory role of CLA-1 in axon regeneration can be genetically uncoupled from synapse function, which explains why knocking down CLA-1 can both promote axonal repair and enhance functional recovery.

### Active zone proteins employ multiple independent strategies to regulate axon regeneration

Why and how active zone proteins, which are typically thought to function in the segment of the axon that does not regenerate, are perplexing questions. Not only do multiple core active zone proteins inhibit regeneration, they also inhibit regeneration across animals. For example, our finding that UNC-13 inhibits *C. elegans* GABAergic axon regeneration is consistent with the role of mammalian uncoordinated-13 (Munc13) in axon regeneration following spinal cord injury (Figure 2B,C) (Hilton et al. 2022). The strong additive phenotype of axon regeneration in *cla-1(-); unc-13(-)* double mutants, along with the subsequent finding that PTRN-1 function is required for enhanced axon regeneration in *cla-1* mutants but not *unc-13* mutants, suggests CLA-1 and UNC-13 inhibit axon regeneration through independent mechanisms (Figure 2B,C, Figure 3C,D). Therefore, presynaptic proteins have developed multiple approaches to inhibit regeneration across species, which suggests there must be a broad mechanistic advantage to inhibiting regeneration with these proteins (Figure 3E).

A comparison of our results with previous data reveals synaptic proteins also regulate axon regeneration in a cell-type specific manner. Here, we found active zone proteins including CLA-1/Piccolo, UNC-10/RIM and UNC-13/Munc13 negatively regulate axon regeneration in GABAergic motor neurons (Figure 2B,C), while a large-scale screen previously revealed that CLA-1 positively regulates axon regeneration in *C. elegans* PLM mechanosensory neurons (Chen et al. 2011). In the same screen, most synaptic vesicle exocytosis genes, including *unc-10/Rim* and *unc-13/mUnc13* were dispensable for PLM axon regeneration. A caveat to this comparison is that hypomorphic mutant alleles of *unc-10* and *unc-13* were used in the large-scale screen of mechanosensory axon regeneration and null mutant alleles were used in this analysis of motor axon regeneration. Future systematic studies will be needed to identify the extent of CLA-1/Piccolo function during axon regeneration in other neuron types and animals, and what determines their context-specific function during axon regeneration.

### CLA-1M, a previously uncharacterized isoform of CLA-1, is sufficient to inhibit axon regeneration cell-autonomously

From a domain analysis of CLA-1, our data reveal that the relatively uncharacterized medium isoform of CLA-1 inhibits motor axon regeneration (Figure 1G,H). Previous studies identified both selective and overlapping functions of CLA-1L and CLA-1S during active zone assembly, synaptic vesicle clustering and presynaptic autophagy during development (Xuan et al. 2017; Xuan et al. 2023; Krout et al. 2023). However, because of the genetic structure of CLA-1, specifically mutating the CLA-1M isoforms to ascertain their individual contributions to synapse development and function has been an unmet goal of the CLA-1 field. In the context of axonal injury, the N-terminal repetitive region of CLA-1, which is specific to CLA-1L, is dispensable for axon regeneration (Figures 1B,D,E). Instead, the in-frame deletion *ok937*, which disrupts CLA-1L and CLA-1M but not CLA-1S, significantly enhanced axon regeneration to the same extent as the putative null allele *ok2285* (Figure 1D,E). On their own, these data suggest three possibilities: that CLA-1L depends on the deleted region for its function; that CLA1M is selectively required; or that CLA-1M and CLA-1L act redundantly to inhibit axon regeneration. Although we are not able to conclusively tease these possibilities apart by specifically disrupting CLA-1M, our finding that expression of CLA-1M in GABA motor neurons is sufficient to inhibit axon regeneration in an otherwise *cla-1*(null) animal reveals a novel, cell-autonomous and potentially specific function of the medium isoform.

### CLA-1 inhibits axon regeneration with microtubule minus-end binding protein PTRN-1

The finding that microtubule minus end binding protein PTRN-1 mediates axon regeneration in *cla-1(-)* mutants, but not in *unc-13(-)* mutants reveals a surprisingly specific interaction (Figure 3C,D). PTRN-1 is critical for maintaining microtubule structure and cargo transport of various molecules and organelles, including active zone proteins and mitochondria, in developing and injured neurons (Marcette, Chen, and Nonet 2014; Richardson et al. 2014; Chuang et al. 2014; Balseiro-Gómez et al. 2022). While PTRN-1 is required to localize CLA-1 during development (Figure 4C,D), our results suggest that CLA-1 has a reciprocal influence on PTRN-1 that modulates microtubule stability or function in injured axons (Figure 5). Our data demonstrate that in wild type axons, injury triggers slow and retrograde-biased trafficking of synaptic protein RAB-3 along the axon commissure that remains attached to the cell body (Figure 5C,D). This phenotype suggests cargo trafficking to and from the site of injury may be congested, potentially because of an increase in cargo or a need to transfer cargo more frequently between dynamic microtubules in an injured axon (Blanquie and Bradke 2018). Our genetic interaction data indicate that in *cla-1(ok937)* mutants, *ptrn-1* function is upregulated (Figure 3). Together with the finding that the speed of RAB-3 trafficking is maintained at its pre-injury level in both directions (Figure 5C,D), and that loss of CLA-1 protects against colchicine-induced microtubule instability (Figure 5G,H), our data support a model where increased *ptrn-1* function in *cla-1(-)* mutants might enhance or speed communication between the site of injury and the cell body as well as transport of regeneration-associated molecules to the injury site.

The specific genetic interaction between CLA-1 and PTRN-1 during axon regeneration raises several interesting questions. Of primary importance is understanding how CLA-1 inhibits PTRN-1 function. The two proteins have recently been reported to physically interact or be localized in close proximity to one another(Sanchez et al. 2021), supporting a model where CLA-1 inhibits PTRN-1 function by physically sequestering or inhibiting a critical domain. An indirect interaction involving transcription regulation, protein expression or post-translational modification is equally possible. It also remains to be determined whether the inhibitory role of CLA-1/Fife/Piccolo during axon regeneration is conserved across different species. The answers to each of the above are critical to further dissecting the roles of CLA-1/Fife/Piccolo and PTRN-1/Patronin/CAMSAP in axon regeneration and may provide additional strategies to improve axon regeneration.

In summary, our findings reveal that knocking down the function of active zone protein CLA-1/Piccolo promotes axon regeneration without disrupting synaptic function and does so by promoting microtubule minus-end binding protein PTRN-1/CAMSAP function and stabilizing microtubule trafficking during the injury response. These data reveal a novel role for CLA-1 and reveal that synaptic proteins have developed multiple independent mechanisms to inhibit axon regeneration.

## Acknowledgments

We thank all members of the Byrne laboratory, Dr. Michael Francis, Dr. Shankar Ramachandran, Dr. Yang Xiang, Dr. Mark Alkema, and Dr. Amy Walker for their experimental suggestions, insight, and critical reading of the manuscript. We thank the *C. elegans* Gene Knockout Consortium (Mitani laboratory and Oklahoma Gene Knockout Consortium), the Caenorhabditis Genetics Center (CGC), which is funded by the NIH Office of Research Infrastructure Program (P40 OD010440), for providing strains and WormBase. We thank the Alkema (UMass Chan Medical School), Francis (UMass Chan Medical School) and Shen (Stanford) laboratories for plasmids and strains.

## Author contributions

W.H contributed to conceptualization, investigation, interpretation, and manuscript preparation. E.M. contributed to investigation, interpretation, and manuscript preparation. A.B.B contributed to conceptualization, interpretation, and manuscript preparation.

## Data and materials availability

All data is available in the main text or the supplementary materials. Strains will be made available upon request.

## Materials and Methods

### Strains

Animals were maintained at 20°C on NGM plates containing OP50 *E. coli* according to standard methods (Brenner 1974). Strains listed below were kindly provided by the Alkema, Francis, Hammarlund and Shen laboratories, and the *Caenorhabditis* Genetics Center, which is funded by the NIH Office of Research Infrastructure Programs (P40 OD010440). To visualize GABA motor neurons, strains were crossed into EG1285 (*oxIs12[unc-47p::GFP, lin-15(+)]*) or IZ629 (*ufIs34[unc-47p::mCherry])*. Mutants and translational reporter used in this study: *cla-1(ok560), cla-1(gk352), cla-1(ok937), cla-1(ok2285), elks-1(ok2762), unc-10(md1117), unc-13(s69), ptrn-1(tm5597), dlk-1(ju476), cebp-1(tm2807), cla-1(wy1233[cla-1::FLPon::mScarlet-I]), cla-1(wy1418[cla-1::GFP])* and *rab-3(wy1332[FLPon::mScarlet-I::rab-3])*.

### Molecular Cloning

Plasmids were built with MultiSite Gateway Technology (Invitrogen) or Gibson isothermal assembly (New England Biolabs) and confirmed by restriction digestion and/or sequencing. Briefly, we cloned CLA-1S/CLA-1E cDNA from pSR58*[Punc-17β::GFPnovo2::CLA-1E::unc-54 3’UTR],* which was kindly gifted from the Michael Francis laboratory at UMass Chan Medical School, into [1-2] CLA-1S entry vector. A proportion of CLA-1M/CLA-1D cDNA was synthesized by Twist Bioscience Inc. and cloned into [1-2] CLA-1S entry vector to make [1-2] CLA-1M entry vector via Gibson isothermal assembly. We also built [1-2] CLA-1MΔPDZ+C_2_ entry vector from [1-2] CLA-1M entry vector via appropriate Gibson cloning primers. For GABA motor neuron-specific expression of CLA-1 isoforms, [4-1] entry vector containing the *unc-47* promoter sequence, [1-2] entry vector containing CLA-1 isoform cDNA and [2-3] entry vector containing *let-858 3’UTR* sequence were recombined into the destination vector pCFJ150 containing *ttTi5605* Mos1 insertion sites and *cb-unc-119(+)* rescue fragment.

### Mos1-mediated single copy insertion (MosSCI)

MosSCI insertion was conducted as previously described (Frøkjaer-Jensen et al. 2008). Briefly, injection was made into EG6699 (*ttTi5605 II; unc-119(ed3) III*) animals following the standard microinjection procedure (Mello et al., 1991). Injection mix included 50 ng/μL pCFJ601(*Peft-3::transposase*), 50 ng/μL plasmids with GABA-specific expression of CLA-1 isoforms, 10 ng/μL pMA122 *(Phsp::peel-1)*, and 2.5 ng/μL pCFJ90 (*Pmyo-2::mCherry*). Injected animals were maintained at 25 °C until starvation. Starved plates were heat-shocked at 34 °C for 4 hours. Afterwards, free-moving animals from the heat-shocked plate were individually transferred into fresh plates. MosSCI insertion was verified by PCR genotyping and absence of fluorescent co-injection marker pCFJ90. Transgenic line was outcrossed to remove *unc-119(ed3)* mutation before crossing into *cla-1(ok2285);oxIs12* for tissue-specific rescue of CLA-1 isoforms.

### FLP/FRT recombination

We used the tissue-specific FLP-on recombination strategy to visualize endogenous CLA-1 and RAB-3 expressions in selected neuron types (Davis et al. 2008; McDonald, Fetter, and Shen 2020). Briefly, we injected plasmid pJR23 *[Pflp-13::NLS::FLP(D5)::NLS::SL2::TagBFP::let-858 3’UTR]* into wild-type Bristol N2 strain at 50 ng/µL with a co-injection marker *Pmyo-2::GFP*. pJR23 plasmid was kindly gifted from the Michael Francis laboratory at UMass Chan Medical School. We then performed UV integration with Stratagene UV Stratalinker 1800 (254nm) and generated integrated transgenic allele *bamIs66[Pflp-13::NLS::FLP(D5)::NLS::SL2::TagBFP::let-858 3’UTR; Pmyo-2::GFP*] following the standard protocol (Evans 2006). We outcrossed *bamIs66* four times before crossing into *cla-1(wy1233[cla-1::FLPon::mScarlet-I]),* or *rab-3(wy1332[FLPon::mScarlet-I::rab-3])*.

### Laser axotomy and quantification

Axotomy experiments were performed as described (Byrne, Edwards, and Hammarlund 2011). Briefly, L4 stage animals were immobilized on 3% agarose pad with 1:12 polystyrene beads. Animals were visualized with a Nikon Eclipse 80i microscope, 100x Plan Apo VC lens (1.4 NA), Andor Zyla sCMOS camera and a Leica EL6000 light source. GABAergic motor neurons were severed at the midline using a 435nm nitrogen pulsed MicroPoint laser at 20 Hz firing frequency. Axotomized animals were recovered in M9 buffer and transferred to fresh OP50-seeded NGM plates. To control technical variability, axotomy experiments were carried out with same-day wild-type controls and appropriate mutant animals. Post-axotomy images were acquired after 24 hours with an Andor Zyla sCMOS camera on a Nikon NI-SSR-930959 microscope and NIS-Elements AR5.02.00 software, or a Perkin Elmer Precisely UltraVIEW VoX confocal imaging system mounted on a Zeiss AXIO (imager.M2) microscope and Volocity 6.3 software. Axotomized commissures were scored as regeneration or not based on the presence of growth cone, and regeneration percentage was determined by the proportion of injured axons that regenerated towards the dorsal nerve cord in each genotype. We also measured the relative length of axotomized commissure by normalizing the length of injured axons to the width of the animals. For example, regenerated axons that reached the dorsal nerve cord would have a relative axon length of 1, whereas axons that did not regenerate typically had a relative axon length below 0.3.

### Nose-touch reversal assay

We measured stimulated reversal movements in axotomized animals 1 hour and 24 hours post-axotomy to assess functional recovery as previously described (Yanik et al. 2004; Byrne et al. 2016). Briefly, 15 out of 16 GABAergic motor commissures, excluding the one in the head, were severed at midlines. Age-matched mock animals underwent the same immobilization process as axotomized animals, except without laser injury. Functional recovery was measured by the number of reversal body bends mock or axotomized animal made following a gentle nose touch from an eye-lash pick. Axotomized animals with no reversal response were referred as “shrinkers.”

### Confocal microscopy imaging and puncta analysis

We acquired FLP-on mScarlet::RAB-3 or CLA-1::FLP-on mScarlet signals in DD neurons with Perkin Elmer Precisely UltraVIEW VoX confocal imaging system mounted on a ZEISS AXIO (imager.M2) microscope using Volocity 6.3 software. Region of interest along the dorsal nerve cord was straightened in Fiji. RAB-3 or CLA-1 puncta number was calculated in Fiji with proper background subtraction, thresholding, binary watershed and analyze particle function. Absence of CLA-1 expressions in cell bodies and commissures of injured axons was individually confirmed by 3D reconstruction in Fiji.

### Live imaging of RAB-3 trafficking and analysis

Time-lapse imaging of RAB-3 in DD3 commissures was performed as previously described with a few modifications (Oliver et al. 2022). Briefly, axotomized or mock control animals were maintained on OP50-seeded NGM plates for 8 hours before time-lapse imaging. For time-lapse imaging, animals were transferred to 3% agarose pad and immobilized with 100mM 2,3-butanedione monoxime (BDM) in M9 buffer. Time-lapse recordings were made on Olympus BX51WI upright microscope with an Olympus UplanXApo 100x/1.45 oil objective through a Yokogawa CSU-X1 Confocal Scanner attached to an Andor iXon Life EMCCD camera at 5 Hz for 30 seconds. Image acquisition was completed using VisiView software by Visitiron Systems. All recordings focused on DD3 axonal commissures and same imaging conditions were applied during three replicates. RAB-3 trafficking kymograph was generated in Fiji to calculate the speed and direction of moving RAB-3 puncta in mock control and injured DD3 axonal commissures.

### Colchicine treatment

Colchicine powder (C_22_H_25_NO_6_, purchased from Thermo Fisher Chemicals) was directly supplemented into NGM plates to reach the final concentration of 0.5mM as previously described (Chalfie and Thomson 1982; Richardson et al. 2014). Animals were raised on 0.5mM colchicine NGM plates seeded with OP50, and their progenies were scored for their GABA motor commissure morphologies or used for mechanosensory assays.

### Quantification of GABA motor commissure morphology

Animals were raised on 0.5mM colchicine NGM plates seeded with OP50, and their progenies were scored for their GABA motor commissure morphologies or used for mechanosensory assays. Animals raised on 0.5mM colchicine NGM plates were immobilized on 3% agarose pads with 300mM sodium azide. We described morphologies of GABAergic motor commissures into the following categories: intact, ectopic branching, and broken. We reported the percentage of animals exhibiting aberrant GABAergic motor commissure morphologies (ectopic branching, broken, and both) under 0.5mM colchicine conditions.

### Mechanosensory assay

We measured mechanosensory response in control and 0.5mM colchicine-treated animals as previously described (Hobert et al. 1999; Richardson et al. 2014). Briefly, touch stimulus was applied alternatively between the head and tail of each animal for 5 rounds. We monitored whether animals responded to each touch by moving away from the stimulus, and we reported the percentage of touch responses each animal displayed with control or 0.5mM colchicine treatment.

### Graphing and statistical analysis

Statistical analysis was performed with Prism (GraphPad). For categorical data (percentage regeneration or percentage morphological defects in GABA motor commissures): data from repeated assays comparing a mutant to corresponding same-day wild-type or mutant controls were pooled with n ≥ 30. Bars represent 95% confidence intervals and statistical significance was determined with Fisher exact test, where *p≤0.05, **p≤0.01, ***p≤0.001 and ***p≤0.0001. For non-parametric data (relative axon length): data from repeated assays comparing a mutant to corresponding same-day wild-type or mutant controls were pooled with n ≥ 30. Bars indicate mean and statistical significance was determined with Kruskal-Wallis test with Dunn’s multiple comparisons test, where *p≤0.05, **p≤0.01, ***p≤0.001 and ***p≤0.0001. Parametric data are plotted as violin plots with medians and interquartile ranges indicated as dashed lines. Significance was calculated with one-way ANOVA with Dunnett’s multiple comparisons test, where *p≤0.05. Parametric data with two independent variables, significance was calculated with two-way ANOVA with Šídák’s multiple comparisons test, where *p≤0.05, **p≤0.01, ***p≤0.001 and ***p≤0.0001.

**Supplementary Figure 1.**
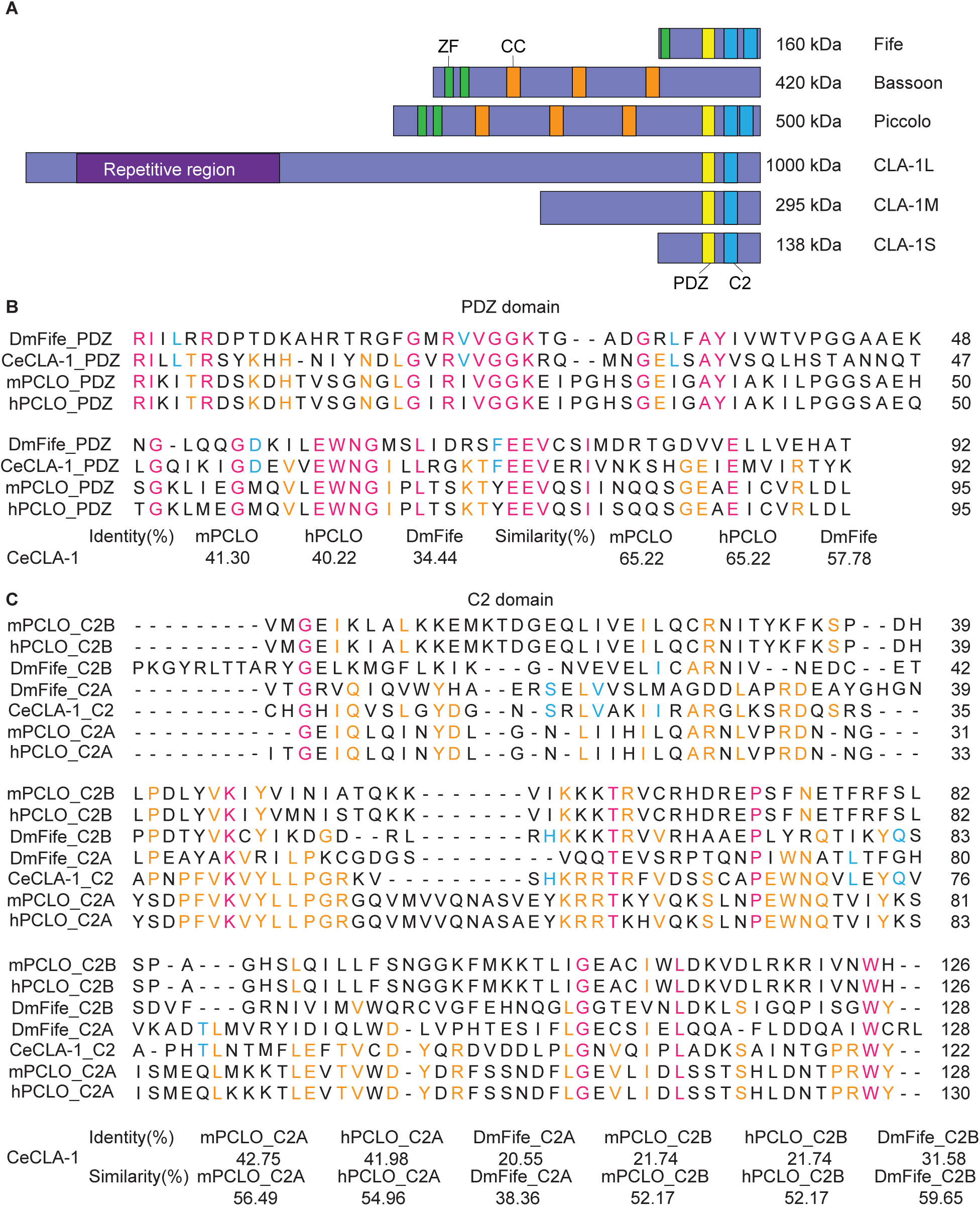
*C. elegans* CLA-1 protein conservation. (**A**) Illustration of CLA-1 and its orthologs: Fife in *Drosophila melanogaster* and Piccolo and Bassoon in mice and human. CLA-1 contains a unique N-terminal repetitive region and two highly conserved C-terminal PDZ and C_2_ domains. Fife and Piccolo possess additional domains, including zinc-finger (ZF) domain and coiled-coil (CC) domain. (**B**) Multiple sequence alignment of PDZ domain among *C. elegans* CLA-1, *D. melanogaster* Fife, mice Piccolo and human Piccolo with Uniprot. Identity and similarity percentage are calculated by pairwise comparison in Protein BLAST. (**C**) Multiple sequence alignment of C_2_ domain among *C. elegans* CLA-1, *D. melanogaster* Fife, mice Piccolo and human Piccolo with Uniprot. Identity and similarity percentage are calculated by pairwise comparison in Protein BLAST. CLA-1 C_2_ domain shares a higher similarity to the C_2_A domain in Fife and Piccolo compared to the C_2_B domain.

**Supplementary Figure 2.**
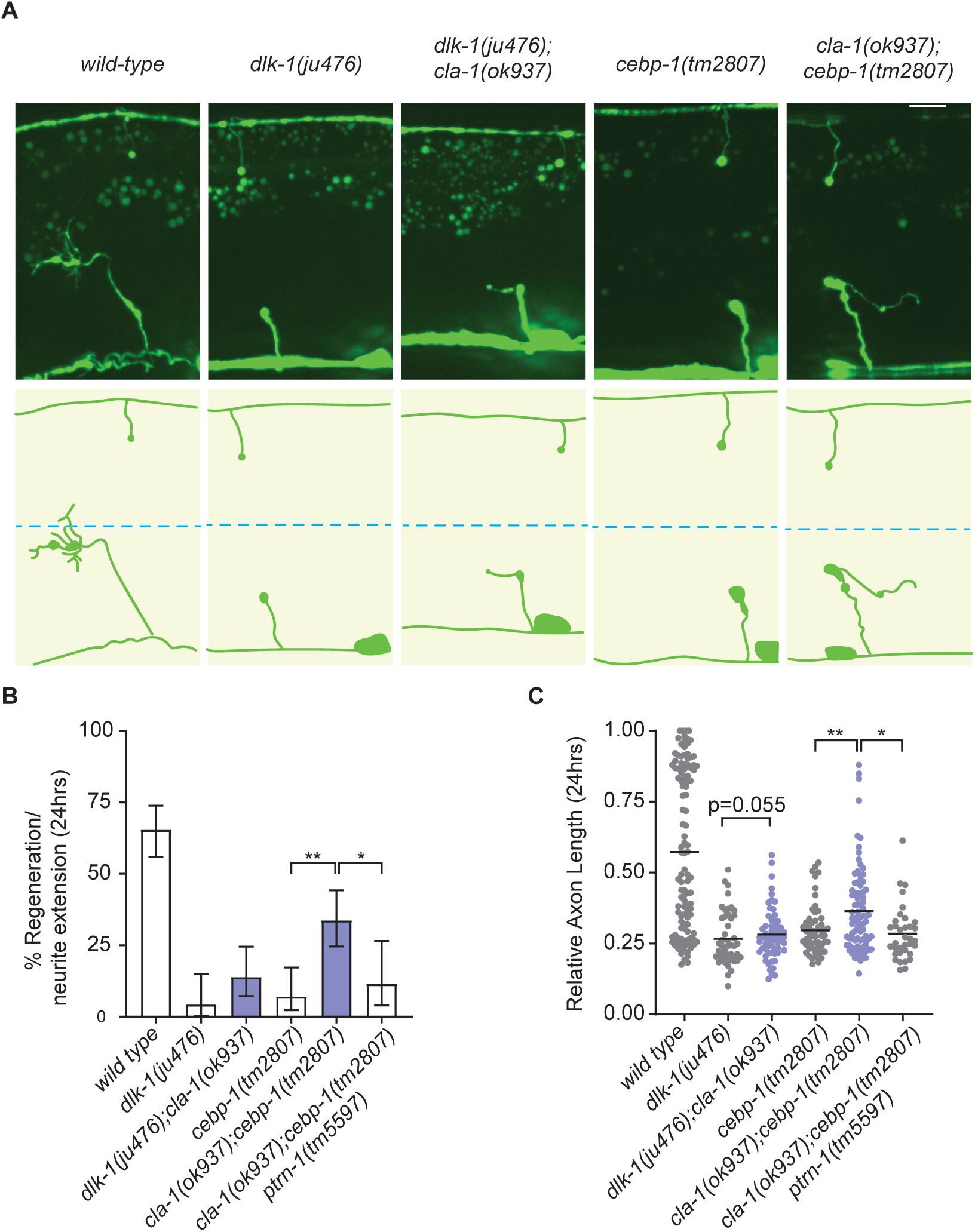
CLA-1 inhibits axon regeneration partially independently of the DLK-1/CEBP-1 pathway. **(A)** Representative regenerating axons in wild-type, *dlk-1(ju476), dlk-1(ju476);cla-1(ok937), cebp-1(tm2807),* and *cla-1(ok937);cebp-1(tm2807)* animals. Blue dashed line represents the dorsal-ventral midline. Scale bar: 10 µm. (**B and C**) Animals lacking components in the p38 mitogen-activated protein kinase (MAPK) signaling, such as DLK-1 and CEBP-1, do not regeneration. Interestingly, loss of CLA-1 promotes filopodia formation at the injured axonal tip in the absence of CEBP-1, but to a less effect in the *dlk-1(ju476)* background. Furthermore, the enhanced regeneration in *cla-1(ok937);cebp-1(tm2807)* is suppressed by the loss of PTRN-1, suggesting that CLA-1 inhibits axon regeneration partially independently or in parallel to the DLK-1/CEBP-1 pathway. N = 104, 47, 65, 57, 86, 35. (**B**) Significance of frequency of regeneration relative to *cla-1(ok937);cebp-1(tm2897)* animals is indicated by *P < 0.05, **P<0.01, Fisher exact test. Error bars indicate 95% confidence intervals. (**C**) Significance of regenerated axon length relative to *dlk-1(ju476);cla-1(ok937)* and *cla-1(ok937);cebp-1(tm2897)* animals is indicated by *P < 0.05, **P < 0.01. Kruskal-Wallis test with Dunn’s multiple comparisons test. Black bar represents median.

**Supplementary Table S1.**

| Sequence ID | Biological Function | Reference |
| --- | --- | --- |
| *elks-1* | Active zone scaffolding protein | Arten et al., 2022 |
| *ptrn-1* | Microtubule minus-end binding protein | Sanchez et al., 2021 |
| *cat-4* | GTP cyclohydrolase I | Lenfant et al., 2010 |
| *cbl-1* | Carbon-sulfur lyase | Lenfant et al., 2010 |
| *dim-1* | Involved in body wall muscle organization. | Lenfant et al., 2010 |
| *eif-6* | Eukaryotic initiation factor | Lenfant et al., 2010 |
| *kel-8* | subunit in the Cul3-RING ubiquitin ligase complex. Involved in ubiquitin-dependent protein degradation. | Lenfant et al., 2010 |
| *nova-1* | mRNA alternative splicing regulator | Lenfant et al., 2010 |
| *pnp-1* | Purine nucleoside phosphorylase | Lenfant et al., 2010 |
| *T19B4.5* | Unknown. Expression in ABalapaaaa, ABalappapa, ABalppappa, germ line and oxygen sensory neurons | Lenfant et al., 2010 |
| *ZC15.3* | Unknown. Expression in head mesodermal cell and neurons | Lenfant et al., 2010 |

## References

1. Apostolakou, A. E., X. K. Sula, K. C. Nastou, G. I. Nasi, and V. A. Iconomidou. 2021. ‘Exploring the conservation of Alzheimer-related pathways between H. sapiens and C. elegans: a network alignment approach’, Sci Rep, 11: 4572.

2. Artan, M., M. Hartl, W. Chen, and M. de Bono. 2022. ‘Depletion of endogenously biotinylated carboxylases enhances the sensitivity of TurboID-mediated proximity labeling in Caenorhabditis elegans’, J Biol Chem, 298: 102343.

3. Balseiro-Gómez, S., J. Park, Y. Yue, C. Ding, L. Shao, S. Ҫetinkaya, C. Kuzoian, M. Hammarlund, K. J. Verhey, and S. Yogev. 2022. ‘Neurexin and frizzled intercept axonal transport at microtubule minus ends to control synapse formation’, Dev Cell, 57: 1802–16.e4.

4. Belew, M. Y., W. Huang, J. T. Florman, M. J. Alkema, and A. B. Byrne. 2023. ‘PARP knockdown promotes synapse reformation after axon injury’, bioRxiv.

5. Blanquie, O., and F. Bradke. 2018. ‘Cytoskeleton dynamics in axon regeneration’, Curr Opin Neurobiol, 51: 60–69.

6. Brenner, S. 1974. ‘The genetics of Caenorhabditis elegans’, Genetics, 77: 71–94.

7. Byrne, A. B., T. J. Edwards, and M. Hammarlund. 2011. ‘In vivo laser axotomy in C. elegans’, J Vis Exp.

8. Byrne, A. B., R. D. McWhirter, Y. Sekine, S. M. Strittmatter, D. M. Miller, and M. Hammarlund. 2016. ‘Inhibiting poly(ADP-ribosylation) improves axon regeneration’, Elife, 5.

9. Byrne, A. B., T. Walradt, K. E. Gardner, A. Hubbert, V. Reinke, and M. Hammarlund. 2014. ‘Insulin/IGF1 signaling inhibits age-dependent axon regeneration’, Neuron, 81: 561–73.

10. Chalfie, M., and J. N. Thomson. 1982. ‘Structural and functional diversity in the neuronal microtubules of Caenorhabditis elegans’, J Cell Biol, 93: 15–23.

11. Chen, L., Z. Wang, A. Ghosh-Roy, T. Hubert, D. Yan, S. O’Rourke, B. Bowerman, Z. Wu, Y. Jin, and A. D. Chisholm. 2011. ‘Axon regeneration pathways identified by systematic genetic screening in C. elegans’, Neuron, 71: 1043–57.

12. Chuang, M., A. Goncharov, S. Wang, K. Oegema, Y. Jin, and A. D. Chisholm. 2014. ‘The microtubule minus-end-binding protein patronin/PTRN-1 is required for axon regeneration in C. elegans’, Cell Rep, 9: 874–83.

13. Czech, V. L., L. C O’Connor, B. Philippon, E. Norman, and A. B. Byrne. 2023. ’TIR-1/SARM1 inhibits axon regeneration and promotes axon degeneration’, Elife, 12.

14. Davis, M. W., J. J. Morton, D. Carroll, and E. M. Jorgensen. 2008. ‘Gene activation using FLP recombinase in C. elegans’, PLoS Genet, 4: e1000028.

15. Ding, C., and M. Hammarlund. 2018. ‘Aberrant information transfer interferes with functional axon regeneration’, Elife, 7.

16. El Bejjani, R., and M. Hammarlund. 2012. ‘Notch signaling inhibits axon regeneration’, Neuron, 73: 268–78.

17. Evans, Thomas C. 2006. ‘Transformation and microinjection.’, WormBook: The Online Review of C. elegans Biology [Internet]. https://www.ncbi.nlm.nih.gov/books/NBK19648/.

18. Frøkjaer-Jensen, C., M. W. Davis, C. E. Hopkins, B. J. Newman, J. M. Thummel, S. P. Olesen, M. Grunnet, and E. M. Jorgensen. 2008. ‘Single-copy insertion of transgenes in Caenorhabditis elegans’, Nat Genet, 40: 1375–83.

19. Gabel, C. V., F. Antoine, C. F. Chuang, A. D. Samuel, and C. Chang. 2008. ‘Distinct cellular and molecular mechanisms mediate initial axon development and adult-stage axon regeneration in C. elegans’, Development, 135: 1129–36.

20. Ghosh-Roy, A., Z. Wu, A. Goncharov, Y. Jin, and A. D. Chisholm. 2010. ‘Calcium and cyclic AMP promote axonal regeneration in Caenorhabditis elegans and require DLK-1 kinase’, J Neurosci, 30: 3175–83.

21. Hammarlund, M., P. Nix, L. Hauth, E. M. Jorgensen, and M. Bastiani. 2009. ‘Axon regeneration requires a conserved MAP kinase pathway’, Science, 323: 802–6.

22. He, L., L. van Beem, B. Snel, C. C. Hoogenraad, and M. Harterink. 2022. ‘PTRN-1 (CAMSAP) and NOCA-2 (NINEIN) are required for microtubule polarity in Caenorhabditis elegans dendrites’, PLoS Biol, 20: e3001855.

23. Hilton, B. J., A. Husch, B. Schaffran, T. C. Lin, E. R. Burnside, S. Dupraz, M. Schelski, J. Kim, J. A. Müller, S. Schoch, C. Imig, N. Brose, and F. Bradke. 2022. ‘An active vesicle priming machinery suppresses axon regeneration upon adult CNS injury’, Neuron, 110: 51–69.e7.

24. Hobert, O., D. G. Moerman, K. A. Clark, M. C. Beckerle, and G. Ruvkun. 1999. ‘A conserved LIM protein that affects muscular adherens junction integrity and mechanosensory function in Caenorhabditis elegans’, J Cell Biol, 144: 45–57.

25. Kim, K., and C. Li. 2004. ‘Expression and regulation of an FMRFamide-related neuropeptide gene family in Caenorhabditis elegans’, J Comp Neurol, 475: 540–50.

26. Kim, K. W., N. H. Tang, C. A. Piggott, M. G. Andrusiak, S. Park, M. Zhu, N. Kurup, S. J. Cherra, 3rd, Z. Wu, A. D. Chisholm, and Y. Jin. 2018. ‘Expanded genetic screening in Caenorhabditis elegans identifies new regulators and an inhibitory role for NAD(+) in axon regeneration’, Elife, 7.

27. Kittelmann, M., J. Hegermann, A. Goncharov, H. Taru, M. H. Ellisman, J. E. Richmond, Y. Jin, and S. Eimer. 2013. ‘Liprin-α/SYD-2 determines the size of dense projections in presynaptic active zones in C. elegans’, J Cell Biol, 203: 849–63.

28. Koushika, S. P., J. E. Richmond, G. Hadwiger, R. M. Weimer, E. M. Jorgensen, and M. L. Nonet. 2001. ‘A post-docking role for active zone protein Rim’, Nat Neurosci, 4: 997–1005.

29. Krout, M., K. H. Oh, A. Xiong, E. B. Frankel, P. T. Kurshan, H. Kim, and J. E. Richmond. 2023. ‘C. elegans Clarinet/CLA-1 recruits RIMB-1/RIM-binding protein and UNC-13 to orchestrate presynaptic neurotransmitter release’, Proc Natl Acad Sci U S A, 120: e2220856120.

30. Lai, C. H., C. Y. Chou, L. Y Ch’ang, C. S. Liu, and W. Lin. 2000. ’Identification of novel human genes evolutionarily conserved in Caenorhabditis elegans by comparative proteomics’, Genome Res, 10: 703–13.

31. Lenfant, N., J. Polanowska, S. Bamps, S. Omi, J. P. Borg, and J. Reboul. 2010. ‘A genome-wide study of PDZ-domain interactions in C. elegans reveals a high frequency of non-canonical binding’, BMC Genomics, 11: 671.

32. Liu, K., Y. Lu, J. K. Lee, R. Samara, R. Willenberg, I. Sears-Kraxberger, A. Tedeschi, K. K. Park, D. Jin, B. Cai, B. Xu, L. Connolly, O. Steward, B. Zheng, and Z. He. 2010. ‘PTEN deletion enhances the regenerative ability of adult corticospinal neurons’, Nat Neurosci, 13: 1075–81.

33. Marcette, J. D., J. J. Chen, and M. L. Nonet. 2014. ‘The Caenorhabditis elegans microtubule minus-end binding homolog PTRN-1 stabilizes synapses and neurites’, Elife, 3: e01637.

34. McDonald, N. A., R. D. Fetter, and K. Shen. 2020. ‘Assembly of synaptic active zones requires phase separation of scaffold molecules’, Nature, 588: 454–58.

35. McIntire, S. L., E. Jorgensen, J. Kaplan, and H. R. Horvitz. 1993. ‘The GABAergic nervous system of Caenorhabditis elegans’, Nature, 364: 337–41.

36. Mukherjee, K., X. Yang, S. H. Gerber, H. B. Kwon, A. Ho, P. E. Castillo, X. Liu, and T. C. Südhof. 2010. ‘Piccolo and bassoon maintain synaptic vesicle clustering without directly participating in vesicle exocytosis’, Proc Natl Acad Sci U S A, 107: 6504–9.

37. Nix, P., M. Hammarlund, L. Hauth, M. Lachnit, E. M. Jorgensen, and M. Bastiani. 2014. ‘Axon regeneration genes identified by RNAi screening in C. elegans’, J Neurosci, 34: 629–45.

38. Oh, K. H., M. D. Krout, J. E. Richmond, and H. Kim. 2021. ‘UNC-2 CaV2 Channel Localization at Presynaptic Active Zones Depends on UNC-10/RIM and SYD-2/Liprin-α in Caenorhabditis elegans’, J Neurosci, 41: 4782–94.

39. Oliver, D., S. Ramachandran, A. Philbrook, C. M. Lambert, K. C. Q. Nguyen, D. H. Hall, and M. M. Francis. 2022. ‘Kinesin-3 mediated axonal delivery of presynaptic neurexin stabilizes dendritic spines and postsynaptic components’, PLoS Genet, 18: e1010016.

40. Park, K. K., K. Liu, Y. Hu, P. D. Smith, C. Wang, B. Cai, B. Xu, L. Connolly, I. Kramvis, M. Sahin, and Z. He. 2008. ‘Promoting axon regeneration in the adult CNS by modulation of the PTEN/mTOR pathway’, Science, 322: 963–6.

41. Richardson, C. E., K. A. Spilker, J. G. Cueva, J. Perrino, M. B. Goodman, and K. Shen. 2014. ‘PTRN-1, a microtubule minus end-binding CAMSAP homolog, promotes microtubule function in Caenorhabditis elegans neurons’, Elife, 3: e01498.

42. Richmond, J. E., W. S. Davis, and E. M. Jorgensen. 1999. ‘UNC-13 is required for synaptic vesicle fusion in C. elegans’, Nat Neurosci, 2: 959–64.

43. Sanchez, A. D., T. C. Branon, L. E. Cote, A. Papagiannakis, X. Liang, M. A. Pickett, K. Shen, C. Jacobs-Wagner, A. Y. Ting, and J. L. Feldman. 2021. ‘Proximity labeling reveals non-centrosomal microtubule-organizing center components required for microtubule growth and localization’, Curr Biol, 31: 3586–600.e11.

44. Schwartz, M. L., and E. M. Jorgensen. 2016. ‘SapTrap, a Toolkit for High-Throughput CRISPR/Cas9 Gene Modification in Caenorhabditis elegans’, Genetics, 202: 1277–88.

45. Shin, J. E., Y. Cho, B. Beirowski, J. Milbrandt, V. Cavalli, and A. DiAntonio. 2012. ‘Dual leucine zipper kinase is required for retrograde injury signaling and axonal regeneration’, Neuron, 74: 1015–22.

46. Silverman, G. A., C. J. Luke, S. R. Bhatia, O. S. Long, A. C. Vetica, D. H. Perlmutter, and S. C. Pak. 2009. ‘Modeling molecular and cellular aspects of human disease using the nematode Caenorhabditis elegans’, Pediatr Res, 65: 10–8.

47. Song, Y., K. M. Ori-McKenney, Y. Zheng, C. Han, L. Y. Jan, and Y. N. Jan. 2012. ‘Regeneration of Drosophila sensory neuron axons and dendrites is regulated by the Akt pathway involving Pten and microRNA bantam’, Genes Dev, 26: 1612–25.

48. Steward, O., R. B. Goldschmidt, and T. Sutula. 1984. ‘Neurotoxicity of colchicine and other tubulin-binding agents: a selective vulnerability of certain neurons to the disruption of microtubules’, Life Sci, 35: 43–51.

49. White, J. G., E. Southgate, J. N. Thomson, and S. Brenner. 1986. ‘The structure of the nervous system of the nematode Caenorhabditis elegans’, Philos Trans R Soc Lond B Biol Sci, 314: 1–340.

50. Wu, Z., A. Ghosh-Roy, M. F. Yanik, J. Z. Zhang, Y. Jin, and A. D. Chisholm. 2007. ‘Caenorhabditis elegans neuronal regeneration is influenced by life stage, ephrin signaling, and synaptic branching’, Proc Natl Acad Sci U S A, 104: 15132–7.

51. Xiong, X., X. Wang, R. Ewanek, P. Bhat, A. Diantonio, and C. A. Collins. 2010. ‘Protein turnover of the Wallenda/DLK kinase regulates a retrograde response to axonal injury’, J Cell Biol, 191: 211–23.

52. Xuan, Z., L. Manning, J. Nelson, J. E. Richmond, D. A. Colón-Ramos, K. Shen, and P. T. Kurshan. 2017. ‘Clarinet (CLA-1), a novel active zone protein required for synaptic vesicle clustering and release’, Elife, 6.

53. Xuan, Z., S. Yang, B. Clark, S. E. Hill, L. Manning, and D. A. Colón-Ramos. 2023. ‘The active zone protein Clarinet regulates synaptic sorting of ATG-9 and presynaptic autophagy’, PLoS Biol, 21: e3002030.

54. Yan, D., Z. Wu, A. D. Chisholm, and Y. Jin. 2009. ‘The DLK-1 kinase promotes mRNA stability and local translation in C. elegans synapses and axon regeneration’, Cell, 138: 1005–18.

55. Yanik, M. F., H. Cinar, H. N. Cinar, A. D. Chisholm, Y. Jin, and A. Ben-Yakar. 2004. ‘Neurosurgery: functional regeneration after laser axotomy’, Nature, 432: 822.

56. Yogev, S., R. Cooper, R. Fetter, M. Horowitz, and K. Shen. 2016. ‘Microtubule Organization Determines Axonal Transport Dynamics’, Neuron, 92: 449–60.

